# Rat anterior cingulate neurons responsive to rule or strategy changes are modulated by the hippocampal theta rhythm and sharp-wave ripples

**DOI:** 10.1101/2024.01.24.577008

**Authors:** M Khamassi, A Peyrache, K Benchenane, DA Hopkins, N Lebas, V Douchamps, J Droulez, FP Battaglia, SI Wiener

## Abstract

To better understand neural processing during adaptive learning of stimulus-response-reward contingencies, we recorded synchrony of neuronal activity in anterior cingulate cortex (ACC) with hippocampal rhythms in male rats acquiring and switching between spatial and visual discrimination tasks in a Y-maze. ACC population and single unit activity responded shortly after task rule changes, or just before the rats adopted different task strategies. Hippocampal theta oscillations (associated with memory encoding) modulated an elevated proportion of rule-change responsive neurons (70%), but other neurons that were correlated with strategy-change, strategy value, and reward-rate were not. However, hippocampal sharp wave-ripples modulated significantly higher proportions of rule-change, strategy-change and reward-rate responsive cells during post-session sleep but not pre-session sleep. This suggests an underestimated mechanism for hippocampal mismatch and contextual signals to facilitate ACC detection of contingency changes for cognitive flexibility, a function that is attenuated after it is damaged.

## Introduction

Adopting an adaptive behavioral response policy, or strategy, in a novel situation requires observation, deduction and memorization of the rules that govern success as well as failure. Optimizing behavior in a changing world involves continuous selection of appropriate strategies, as well as inhibition of less successful ones. Furthermore, alternative, possibly more beneficial, behavioral strategies should be developed and tested. This all requires tracking outcomes over successive iterations, an application of working memory (Malenka, et al., 2009; Yu and Frank, 2015). This would benefit from neural coding for the presence or absence of rewards, which is found in anterior cingulate cortex (ACC) and upstream structures like the hippocampus (HPC; e.g., Tabuchi et al., 2003; Hyman et al., 2011). Furthermore, the HPC is strongly implicated in spatial and contextual coding, but little is known about how hippocampal signal processing modulates neural representations of behavioral flexibility in ACC. Note that the nomenclature here follows van Heukelum, et al (2020): ACC for medial prefrontal cortex (mPFC), A32 for prelimbic area (PL), A25 for infralimbic area (IL), and A24 for anterior cingulate cortices Cg1 and Cg2, also referred to as dorsal anterior cingulate cortex (ACd). In addition to working memory, ACC and closely associated (Arikuni, et al., 1994) prefrontal cortex, (Goldman-Rakic, 1987, 1995; Lenartowicz and McIntosh, 2005; D’Esposito et al., 1995), mediate other cognitive processes underlying rule-based strategy selection: attention (Shallice, 1988), flexible adaptation of response-selection mechanisms in the presence of novel, fluctuating or ambiguous contexts (Dias et al., 1996; Robbins, 2007), and consolidation of memory (Frankland and Bontempi, 2005). Indeed, patients with prefrontal cortex damage are impaired in non-cued rule-shifting tasks, showing persistence errors (Milner, 1963; Drewe, 1974). Neurophysiological recordings, as well as neuropsychological studies, indicate a crucial role for the rodent ACC and its primate homolog in encoding task-relevant information such as reward (Pratt and Mizumori, 2001), reward rate (Genovesio, et al., 2005), and action-outcome contingencies (Killcross and Coutureau, 2003; Del Arco, et al., 2017). Further, ACC and closely associated areas can represent the current task rule (Mansouri et al., 2006; Rich and Shapiro, 2009; Bissonnette et al., 2013; Chiang et al., 2021, but see Malagon-Vina, et al., 2018) as well as the behavioral strategy adapted to the current task rule (Genovesio et al., 2005; Durstewitz et al., 2010). Moreover, in tasks involving purely spatial strategies, PFC activity has been reported to shift after instatement of a new rule, but before the animal’ s volitional change in strategy (Karlsson, et al., 2012; Powell & Redish, 2016). However, it is unknown whether this result generalizes to cue-guided strategies, whether distinct PFC populations encode distinct types of shifts, and how these behavioral flexibility-related prefrontal activities interact with the HPC during behavior and subsequent sleep.

It is not yet understood how executive and memory functions are linked and coordinated in the HPC-ACC axis. Specifically, how do populations of neurons within the ACC encode different types of task-relevant information in order to both detect sudden rule changes and participate in the learning a new behavioral strategy or switching to a previously acquired one following such changes? It is also unclear whether the same ACC neurons are coordinated with HPC during rule change detection and during initiation of new behavioral strategies. Furthermore, little is known about such cells’ reactivation during sleep in synchrony with hippocampal ripple activity, when memory consolidation would occur (Girardeau, et al., 2009). Computational models of strategy selection in the ACC predict that distinct neuronal populations would subserve functions such as reward detection/processing, goal/strategy selection, and action planning (Hasselmo, 2005; Martinet et al., 2011; Dollé et al., 2018). Other models suggest that off-line replay may play a critical role in making context- or item-selective activity emerge during learning (e.g., Raudies and Hasselmo, 2014).

A possible substrate for memorizing the outcomes of previous actions in order to detect rule changes and implement rewarding strategies could involve synchronization of hippocampal and ACC activity by the theta rhythm, as has been observed during memory tasks (Hyman, et al., 2005; Jones and Wilson, 2005; Siapas, et al., 2005; Benchenane, et al. 2010; cf. Fries, 2015). A mechanism proposed to underlie consolidation of labile hippocampal memory traces into more stable cortical representations invokes the activation of ACC neurons in conjunction with hippocampal oscillations during training sessions. This coordinated activity would then be reactivated in association with hippocampal memory replay during awake and asleep ripples (Jadhav, et al. 2016; Peyrache et al., 2009; Tang, et al., 2017; Yu, et al., 2018). SWR activity is coordinated among brain structures and this is associated with learning (Eschenko, et al., 2008; Ramadan, et al., 2009; Lansink et al., 2009; Girardeau et al., 2009; Roux, et al., 2017; Tatsuno, et al., 2020). Indeed, prefrontal cell assemblies form during periods of high HPC-ACC coherence upon initial acquisition of new rules (Benchenane, et al., 2010) and are reactivated during SWR in subsequent sleep (Peyrache, et al., 2009). Thus, the ACC can be viewed as a neural substrate for learning and evaluating the suitability of action-outcome contingencies (or rules), and orchestrating the execution of appropriate behavioral strategies (Bunge, 2004), benefitting from hippocampal contextual and memory signals.

In order to reveal neuronal activity profiles that could underlie key prefrontal functions, here we examined single unit activity in ACC subregions A32, A24 and A25 in relation to local field potential (LFP) oscillatory activity in the HPC of rats acquiring and shifting between spatial and cue-guided strategies in a Y-maze (original data set at https://crcns.org/data-sets/pfc/pfc-6/). Here we define strategies operationally as descriptions of the patterns of behavioral responses relevant to the structure of the task and the maze. We make no assumptions about the internal representations and neural mechanisms engaged when the animals perform the task. The task was designed to emulate the extradimensional set shifts of the Wisconsin Card Sorting Task employed to diagnose cingulate cortex deficits in human patients (Berg, 1948; Grant et al., 1949; Milner, 1963). Once the rats reached criterion performance in one task, the rule was changed (“RC sessions”). Then, the animal had to learn that the previous strategy was no longer optimal, and had to learn or shift to a new strategy. We examined sessions when the animal changed strategies (“SC sessions”), and these could include strategies not adherent to the current rule. Individual neurons with firing rate changes in these sessions were identified as RC- or SC-responsive. We examined the modulation of these neurons by the hippocampal theta rhythm (8 Hz) during behavior, as well as by HPC sharp wave-ripple oscillations (SWRs) during sleep before and after the training sessions, since these would reflect preferential responsiveness to potentially informative hippocampal signals.

## Results

### Behavior

In the Y-maze, each trial began in the start arm. A central barrier was removed and a cue was lit in the left or right arm in pseudorandom sequence. To obtain a liquid reward, the rat had to go to the end of the appropriate reward arm according to the current rule. In the initial rule, the rewarded arm was on the right side (lit or unlit), and the next reward contingency required visits to the lit side (left or right), then the left side, and then the unlit (“dark”) side (**Fig. 1A**). A different rule was applied (Rule Change: RC) after the rat reached a criterion level of 10 consecutive rewarded trials (or 11 rewards in the previous 12 trials). Learning was by trial and error. This stringent criterion was applied because some rats tended to explore other strategies after initial acquisition and this assured that the performance level had become rather stable.

**Figure 1.**
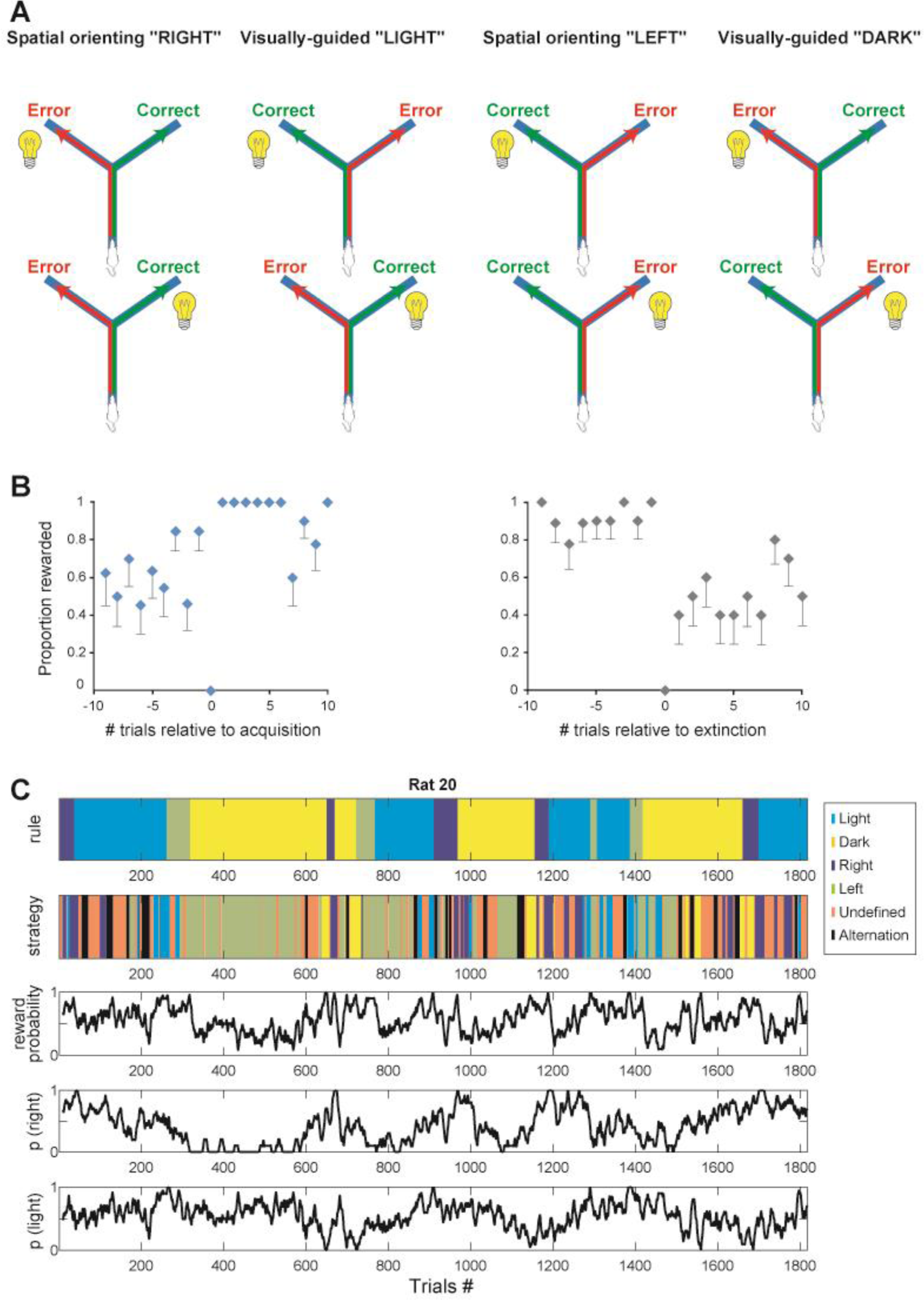
Task and behavior. **A)** Schema of the four Y-maze task rules. A visual cue is lit at the end of one of the two reward arms in a pseudo-random sequence. **B)** (Left) Reward rate changes upon first acquisition of each rule. The first correct trial is at trial zero. Blue diamond=mean; whiskers=SEM. If no whiskers, SEM=0. Data are from 13 sessions of initial rule acquisition of various reward contingencies in the five rats. (Right) Reward rate changes relative to first unrewarded trial revealing change of the task rule (trial number zero). Chance levels of reward after rule change are consistent with initial persistence of the previously rewarded strategy (black diamonds=means; whiskers=SEM). (n=10 sessions). **C)** Behavior in rat 20 over all sessions. Top row: Task rule. Second row: Strategies used by the animal. Third row: Probability of reward. Fourth row: Probability of going to the right arm. Bottom row: Probability of going to the lit arm. Probabilities are calculated over sliding windows of 10 trials.

The five rats performed a total of 3322 trials over 108 sessions. Two rats succeeded at learning only two of the rules (then recordings had to be stopped for technical reasons), and two others learned three, while one rat learned all four, and then continued on to make a total of 17 extradimensional (i.e., between spatial and visual discrimination) strategy switches between the four rewarded rules (e.g., **Fig. 1C**). Learning the first rule (Right) took about 10.2±3.7 (SEM) trials on average, while acquiring the Light rule took much longer, 104.2±50.1 trials. Switching to the Left rule then required 34.3±7.8 trials, and then switching to the Dark rule (only by rat 20) took 319 trials. In most cases, performance did not appear to improve gradually from chance to criterion levels – rather, the animals’ performance levels changed abruptly (**Fig. 1B**; cf., Gallistel et al., 2004). Sometimes animals also spontaneously performed alternation even though it was not rewarded, and this, as well as Right, Light, Left and Dark, are considered as “defined” strategies hereafter. Note, however, that when a Bayesian descriptive model of the behavioral data tested 256 rule combinations (see Supplementary Annex), the animals’ strategies could be described in other frameworks, such as persistence to the same path (left or right) or cued arm (lit or unlit), as well as alternation. We refer to the defined strategies here for simplicity.

The task rule was changed in 24 sessions and, in 16 of these, the rats also made at least one strategy change, that is, extinction (cessation) of the previously rewarded strategy, or adoption of another defined strategy, rewarded or not. After rule changes, 11.4±8.4 trials were required for the animals to abandon the previous strategy, with the exception of one case where a rat persisted for 280 trials before acquiring the next rule. This suggests that, apart from this exception, the rats did not develop habitual behavior from overtraining. Rather this is consistent with them executing goal-oriented navigation strategies, which involve the hippocampus-ACC network (see e.g., Khamassi and Humphries, 2012). For analyses, the criterion of a minimum of six consecutive trials was selected to define a sequence (or “block”) of trials with a given strategy. Simulations of random choices for this behavioral data set showed that this criterion is sufficient to exclude fortuitous inclusion of stochastic choices (see Methods). The animals followed a defined strategy in 170 blocks that included 2424 trials (for example, see **Fig. 1C**). In a total of 898 trials (27% of all trials) the strategy was classified as undefined. The rats consistently adhered to a defined strategy for the entire session in only five sessions. Prior to acquisition of a rewarded task rule, the rats did not tend to simply follow an undefined strategy. Rather, defined strategies other than the currently rewarded one were employed.

The rats changed strategies 118 times, sometimes more than once in a session. Of these 24 were shifts between the task-relevant spatial and visual strategies (Right, Left, Light or Dark). In 55 cases, rats shifted between a defined and an undefined strategy, and they shifted between a task-relevant strategy and the Alternation strategy 39 times. No spontaneous reversals were observed (that is, between Right and Left, or between Light and Dark strategies), nor were they ever rewarded.

In summary, among the 108 recording sessions, these analyses enable us to distinguish the following different types of sessions that are analyzed below:

- 6 *RC-only* sessions, when a rule change (RC) was imposed but the animal made no strategy change (SC) (rather it tended to persist in adhering to the previous rule);
- 65 *SC-only* sessions when a strategy change occurred (not necessarily to the currently rewarded strategy), but there was no RC;
- 18 *RC+SC* sessions, when at least one SC and one RC event occurred; and
- 19 *Training* sessions, when neither a SC nor a RC occurred.

### Electrophysiological recordings

The anatomical distribution of the recording sites of the ACC cells is shown in **Fig. 2**. The largest group was located in layer 5 of A32, although layers 2/3 and 6 of A32 and areas A24 and A25 were also represented.

**Figure 2.**
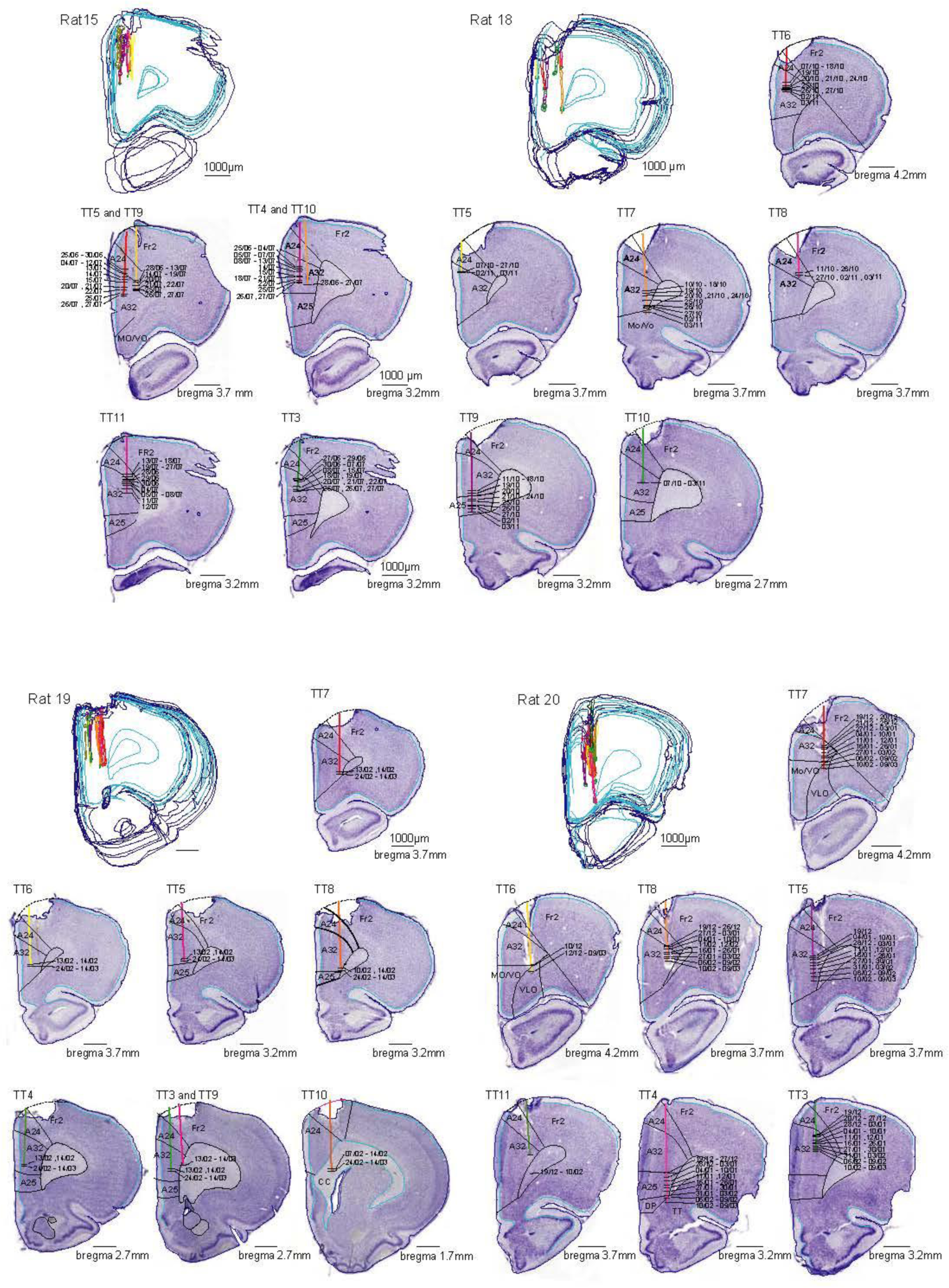
Anatomical localization of recording sites in the rat ACC. For each rat, the image to the upper left is a 3D reconstruction of color-coded tetrode (TT) tracks. The dates are indicated for each recording site. The histology for the fifth rat had technical problems and is not shown.

### Activity transitions at the population level in relation to RC and SC

We first tested if population activity changed relative to RC and SC, as shown previously for criterion performance in rewarded tasks (cited in the Introduction). Population vectors of overall activity of simultaneously recorded neurons were examined for each session to identify those trial(s) when changes in firing rate (“activity transitions”) occurred. These transition points correspond to those trials where there is a major change in the population vector of activity of all recorded neurons. These activity transition trials were then compared to the trials when RC and SC events occurred and the delays were calculated. Specifically, a (z-scored) population vector correlation matrix was constructed with the element at row *i* and column *j* quantifying the correlation between the population vectors at trials *i* and *j* (cf. **Figure 3**). A matrix block decomposition algorithm detected the transition points by searching for the presence of distinct “sub-matrices” within the matrix. These correspond to sequences of trials with distinct patterns of population activity. This process identified and excluded inter-trial comparisons outside the blocks (e.g., in the top row of **Figure 3**, the blue submatrix blocks at the lower left and upper right, called the “off-diagonal” terms) while preserving the maximal amount of information of the matrix (i.e., the highly correlated sub-matrix blocks at the upper left and lower right of Fig. 3). Block decomposition was performed when the cumulated correlation of off-diagonal terms represented less than 1/3 of the total correlation in the matrix. This analysis detected neural activity transitions in 100/108 (92.6%) sessions (see **Fig. 3A**). Of these, 81 sessions contained at least one behavioral event (RC or SC). Interestingly, a population activity transition occurred within four trials before or after a behavioral event (either a RC or a SC) in 68 of the 81 sessions (i.e., 84%; **Fig. 3B**). This is more than expected by chance since a Monte Carlo distribution of 1000 population transitions randomly generated in these sessions yielded only 28% of cases with only four or fewer trials between neural and behavioral events (binomial test, z=11.0, p<.001). Note, however, that there was also a population activity transition in all 19 Training sessions (where no RC or SC event occurred), consistent with previously reported lability of ACC single neuron activity (e.g., Rich and Shapiro, 2009, p. 7214; 12.6% of their neurons; Tanaka, 2007). This could reflect gradual neural processes underlying acquisition of new rules. (We speculate that this corresponds to our subsample of neurons transitioning between attractor states, but this alone may not be sufficient to alter behavior. Rather, this might need to be concerted with some critical number of other neurons, and perhaps in other brain structures as well. Thus, such a population ensemble could develop over the course of several training sessions as more cells acquire the appropriate behavioral correlate and become synchronous, leading to a behavioral change.) Conversely, in 8 (9%) of the 89 sessions where there was an RC or SC behavioral event, no neural activity transition was detected in the respective samples of neurons recorded then.

**Figure 3.**
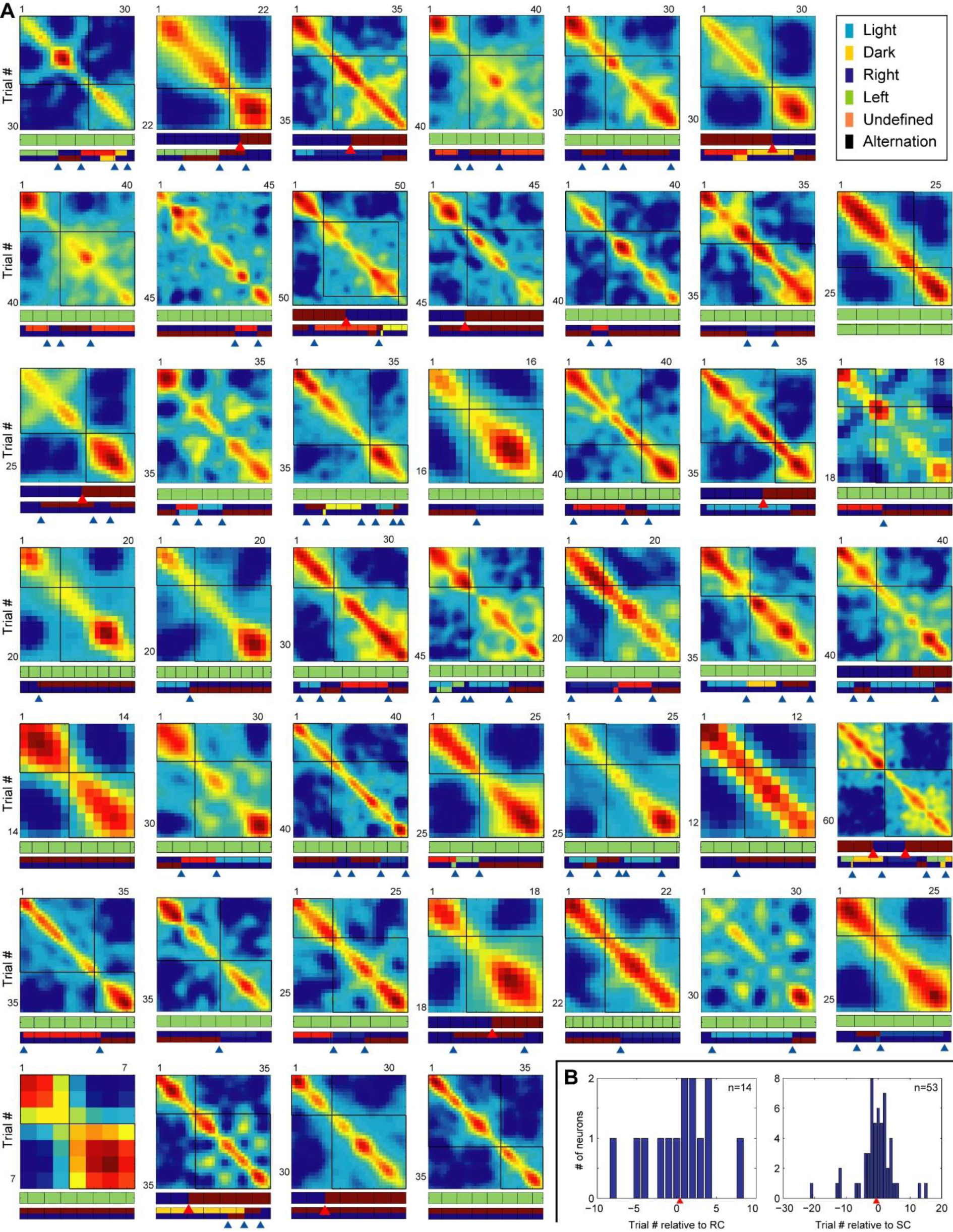
Transitions in population activity over the course of the sessions. **A)** Each box plot shows the correlation matrix of the population vectors between different trials (rows and columns) of an example session. (Hotter colors indicate higher correlations). Crossbars represent the automatically determined decomposition of the matrix into sub-blocks along the diagonal. (Cases with no crossbars are presented to demonstrate how sub-blocks could be absent – in these cases, decomposition led to supra-threshold information loss since off-diagonal terms contained substantial correlations in population activity between trials). Below the matrices, top bars indicate when task rules changed (red arrowheads). Lower bars indicate when strategies changed (blue arrowheads). In the bars, dark blue represents undefined strategies while other colors are defined strategies with an arbitrary color code here. Within each plot, a given color indicates the same rule or strategy. **B)** Incidence of neurons with shifts in population activity at various delays relative to RC or SC behavioral events (which occur at trial zero). Red arrowheads point to means (SC: 0.36 trial; RC: -0.62 trial).

Since multiple behavioral events in the same session could lead to possible confounds, we first examined sessions with only a RC or a SC. In five of the six RC-only sessions a population vector transition occurred 2.3±1.6 trials (median 2.5 trials) *after* the first trial revealing that the RC had occurred. Thus, this neuronal population activity responded rapidly to the RC, even though no explicit cue was presented to signal the RC’s. Strikingly, in the 65 SC-only sessions, the neural activity transition occurred on average 1.0±0.86 trial (median = 1) *before* the first trial of the block of trials qualifying as an SC event. Thus, the transitions in ACC population activity tended to anticipate SCs. Analyses of RC+SC sessions were inconclusive with the population vector approach, and these will be examined further below at the single-cell activity level. In summary, the transitions in mPFC population activity tended to occur in conjunction with RC or SC events.

In order to examine the relation of hippocampal activity with RC and SC responses in ACC, further analyses focused on single neuron activity modulation by hippocampal theta and SWR. First, however, it was necessary to determine if the population responses also appeared in individual neurons. In contrast with the population activity, the individual neuron analyses distinguish firing rate changes at different points of the maze. This permits reward site activity to be set aside, since this could lead to potential confounds related to changes in reward consumption prior to versus after RCs and SCs.

### Firing rate transitions in single ACC neurons in relation to RC or SC

Of the 2290 recorded ACC neurons, 1888 with average firing rates superior to 0.3 Hz in at least one of the task periods (pre-start, post-start, pre-arrival at reward site, post-arrival) were retained for these analyses. Putative interneurons comprised 19% of the cells recorded, and 79% were classified as putative pyramidal cells, while 2% could not be clearly identified. Firstly, 1337 (71%) of the 1888 cells showed behaviorally correlated activity, that is, the firing rate significantly varied across the four trial periods (ANOVA, p<.05). This proportion was similar considering only pyramidal cells as well, and also in the subgroup consisting of A32 layer 5 cells. These proportions are compatible with previous reports of data recorded in animals performing other tasks, employing analyses using different analytical criteria (50%: Pratt and Mizumori, 2001; 62%: Mulder et al., 2003; 46%: Rich and Shapiro, 2009).

To provide an overview, neuronal activity was plotted as a function of trials prior to or after RC and SC events. This revealed numerous neurons that appeared to change firing rate relative to these events (**Figure 4)**. These transitions could be either increases or decreases in firing rate after the RC or SC events, and occurred in cells with high or low firing rates. To more precisely compare the timing of transitions in firing rates of neurons relative to when RC’s and SC’s occurred, Monte Carlo analyses (Fujisawa et al., 2008) examined single unit activity along the maze. This only considered maze regions visited on all trials of the session. This analysis first searched for positions on the maze where actual spike activity exceeded shuffled data first for trials before vs after the 5^th^ trial of the session, then before and after the 6^th^ trial, etc., seeking the trial that had the greatest difference in average firing before vs. after it. These data are represented as the rows of color raster plots of 2^nd^ column of **Figs. 5A-E**. For example, in a session with 40 trials, one of the random shuffles between trials 1 through 5 and trials 6 through 40 could exchange the sequential bins of trial 1 with that of trial 36, trial 2 with trial 13, trial 3 with trial 40, trial 4 with trial 6 and trial 5 with trial 27. The firing rates in the two surrogate groups were compared for each of the spatial bins, then accumulated with data from other such shuffles to create spike density functions for the shuffled data. Then, analogously to transition detection in the population analysis of **Fig. 3**, the greatest “peak transition trials” were determined, and then compared to the timing of the SC or RC trials. The null hypothesis is that the timing of these peak transition trials is unrelated to these behavioral events.

**Figure 4.**
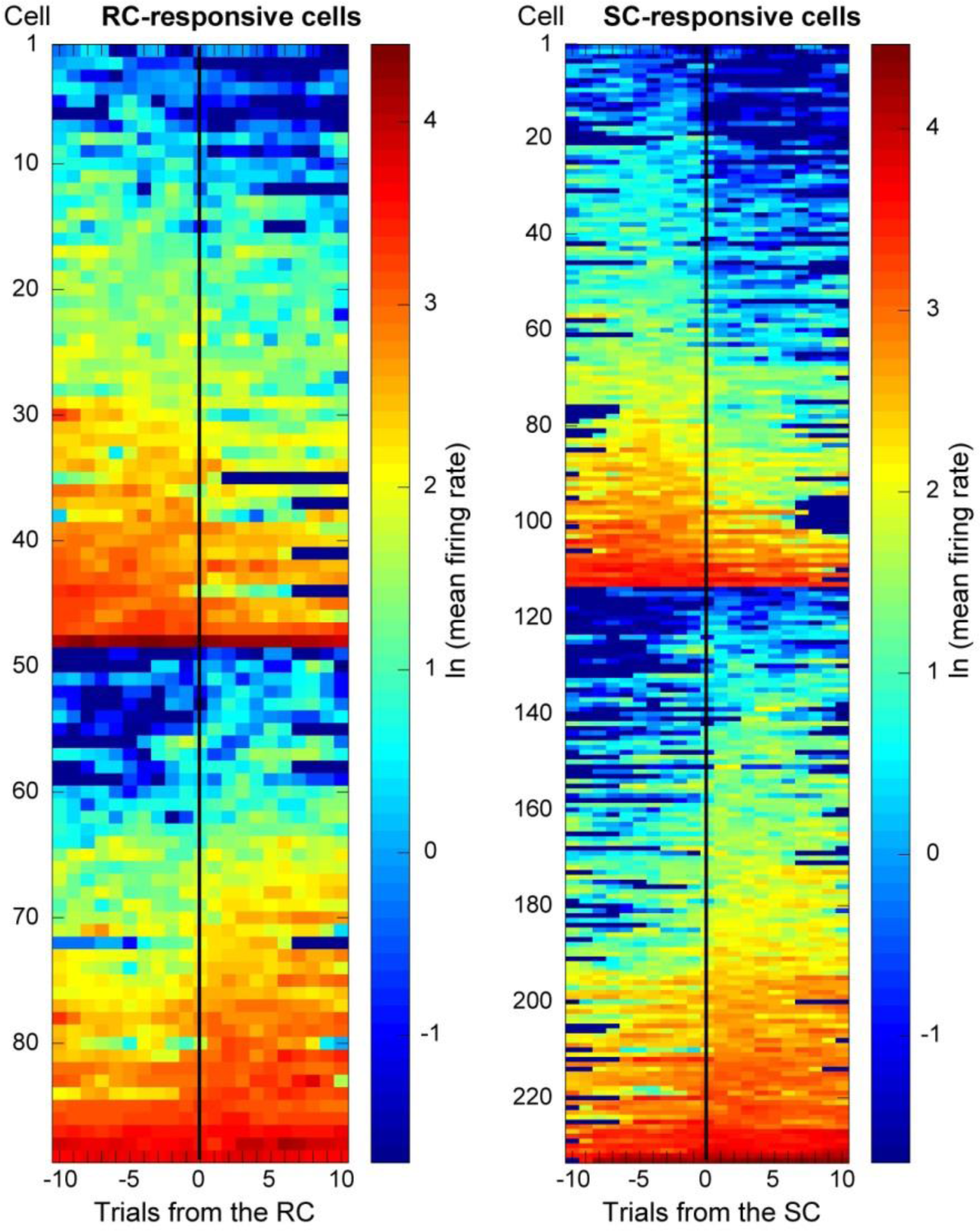
Overview of significant firing rate changes in representative individual neurons before and after RC and SC. Decreases in activity are at the top, and increases at the bottom. The color scale is the natural logarithm of the firing rate. Each row corresponds to one neuron. Plots are synchronized to the trial when the RC could be detected by reward absence, or the trial when the SC occurred.

**Figure 5.**
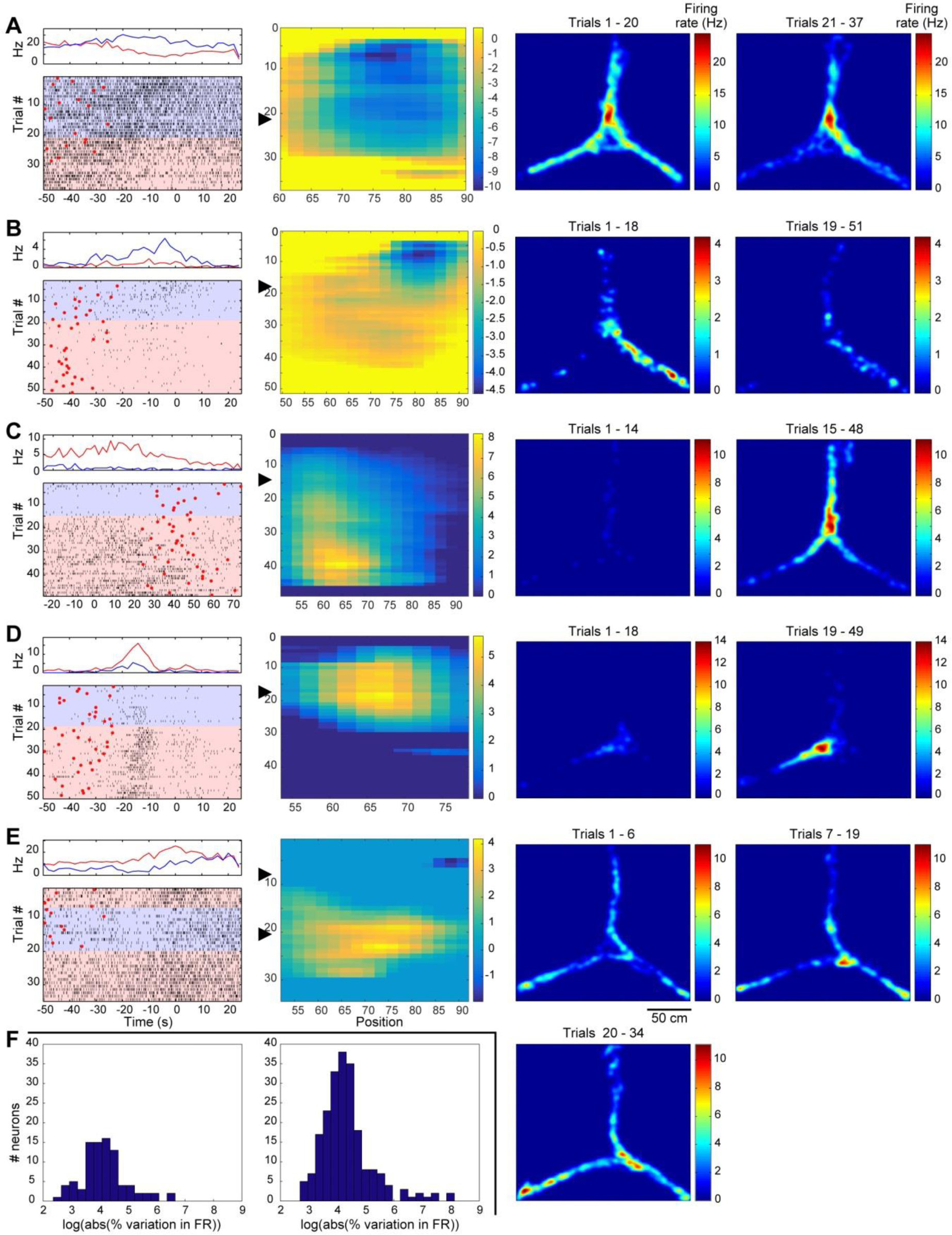
Examples of RC and SC associated changes in firing rate. A through E) Data from representative neurons. The left column contains raster displays and above them, the corresponding histograms of spiking activity, both as functions of time. Blue (or red) histogram traces represent the firing rate before (or after) the RC or SC, corresponding to the color-coded shading of the raster displays. Each row in the rasters is a single trial. In rasters, when the red circles are after zero they correspond to reward site arrival, and zero is the trial start event in these cases. When the red circles are before zero, they indicate the trial start events, and zero is the reward site arrival time. (In E, the histogram for trials 1-6 is not shown.) The plots in the second column show all significant Monte Carlo bootstrap SDF values (trial number on y-axis and linearized position on the maze on the x-axis). Only deep blue rectangles (zero) had non-significant SDFs. The color scale represents the magnitude (in units of spikes/s) of the difference between the SDF value and the upper or lower 2.5% confidence limits. Arrowheads indicate the RC or SC). The third and fourth columns show the spatial distribution of the neuron’s activity before and after the RC or SC. The start arm is above. **A)** A rule-responsive neuron with stable activity at the decision point and a rule-modulated activity in the arrival arms. **B)** A rule-responsive neuron with rule-modulated activity in the left arrival arm. **C)** A strategy-responsive neuron with activity selective for the start arm and decision point. **D)** A strategy-responsive neuron with activity selective for the decision point and right arrival arm. **E)** A strategy-responsive neuron with transitions in activity at two strategy changes on trials #7 and #20 (hence the three spatial plots). The activity is reduced at the decision point while the animal follows the correct Light strategy between trials #7 and #19. The activity remains low during two trials after the task rule change (when the Light strategy is no longer rewarded), but resumes on trial #20 when the rat started a block of Alternation trials. **F)** Distribution of the incidence of neurons with respective variations in firing rate following changes in rule (left) or strategy (right). FR= firing rate. X-axis ranges vary among Figures since they are limited to maze positions the rats occupied on every trial of the respective sessions.

Among the 114 neurons analyzed from the 6 RC-only sessions, 50 (43.9%) showed at least one spike density function (SDF) difference exceeding the shuffled distribution, indicating there was a significant difference in firing rate between at least one of the pairs of blocks of trials in at least one bin on the maze. Similarly, among the 846 neurons analyzed from the 65 SC-only sessions, 400 (47.3%) showed a significant SDF difference (Monte Carlo, p<0.05, two-tailed). **Figs. 5A and B** show data from two representative RC-responsive neurons and **Figs. 5C-E** show three representative SC-responsive neurons. **Fig. 5F** shows the wide range of firing rate increases or decreases in comparisons between blocks before vs. after the RC or SC.

To detect transitions in single neuron activity, the greatest values of these inter-block differences were determined. Then, for each neuron, the incidences of trials separating blocks with these peak SDF values were plotted as a function of the trial when that session’s RC or SC occurred. Since these distributions did not pass the Lilliefors test for normality (p>0.05; **Fig. 6A**), they were fitted with beta distributions to determine normative values. Strikingly, in **Fig. 6B**, the mean delay was 3.9±2.5 trials after the trial when the RC became evident in RC-only sessions (median=3 trials after; significantly different from zero, t-test, p<0.001, n=8649). The mean was also 3.8±2.3 trials after the RC for RC+SC sessions (median=3 trials after, again significantly different from zero, t-test, p<0.001, n=9311).

**Figure 6.**
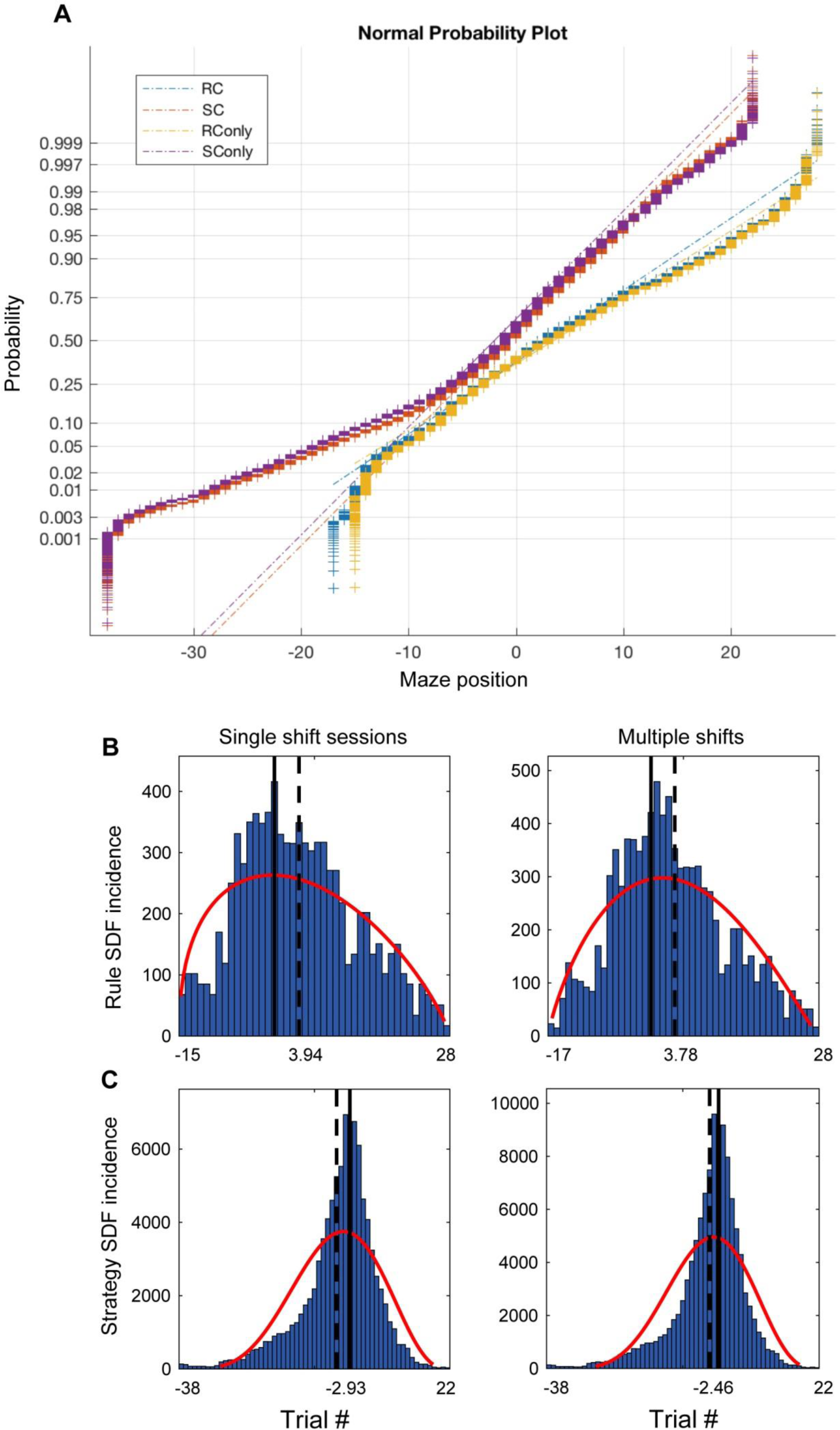
Distributions of significant SDF values for RC-responsive and SC-responsive neurons. **A)** Spatial distributions of significant SDF values do not follow a normal distribution since they deviate from the best fit regression bars (dashed lines). **B and C)** Beta distribution fits (red curves) of incidences of significant SDF values for rule and strategy changes respectively in either Single shift sessions or Multiple shift sessions. Vertical lines represent the trials when the RC (top row) or SC (bottom row) occurred. Vertical dashed lines represent the means of the fitted beta distribution. The SDF distribution means occur after the RCs, but before the SCs.

In SC-only sessions, the mean of the beta distribution of the SDF values was 2.9±1.4 trials *before* the SC event; (**Fig. 6C**; median=1 trial before; significantly different from zero: t-test, p<0.001, N=95629). Similarly, in RC+SC sessions the mean was 2.5±1.3 trials earlier than the SC (median=1 trial before; t-test, p<0.001). Thus, individual ACC neurons can respond rapidly to changes in action-outcome contingencies (i.e., RC’s), or precede adaptive responses to these changes (i.e., SC’s). These results are consistent with those of the population analyses (2.3±1.6 trials after RC and 1.0±0.86 trial before SC).

While it seems counter-intuitive that some transitions occur prior to RC (**Figs. 3B** and **6B**), this may be related to the lability of ACC neurons, or to anticipation of RC by the more experienced rats. The reduced incidence for separations being observed between blocks for early and for late trials could be related to fewer combinatorial samples there. However, the mean values following RC or preceding SC events cannot be due to these potential confounds.

One possible confound for interpreting RC responses in the SD tasks is that the neuronal activity could be correlated with action value, rather than the RC per se. For example, in the Right rule, each trial (rewarded or not) reinforces the value of the appropriate action. This value progresses over trials, but, after RC’s, it decreases for the previous rule, and increments on each trial with the new rule. There is a similar issue with the visual discrimination rules, and thus we extended the concept of action value to “strategy value”. This emulates, in a simple and parsimonious way, the strategy value learning process in relevant computational models (Dollé, et al., 2018; see Supplementary Figure 1). Strikingly, multiple regression analyses showed that the activity of only 8/50 (16%) of RC-responsive neurons was significantly correlated with one of the strategy values (p<.05, significance threshold set with bootstrapping with 10,000 shuffles). Thus, most RC responses cannot simply be a confound with strategy value. Interestingly, firing in 288 other neurons was exclusively correlated with strategy value, and their modulation by hippocampal rhythms is discussed below.

Another possible confound concerned reward rate (RR), which could toggle between 100% and 50% after RCs or SCs. But the firing rate of only 4/50 (8%) RC-responsive neurons was correlated with RR (p<.05, regression analyses with bootstrapping, with rate averaged over six trials). Similarly, only a minority of SC neurons was significantly correlated with RR (16/400, 4%), and thus this was not a major confound. However, 59 neurons were exclusively correlated with RR. RR correlates have been previously observed in ACC (e.g., Genovesio et al., 2005) and would be useful for continuous monitoring of task performance levels, complementing the RC-responses.

### Modulation of neurons by hippocampal SWRs in RC and SC sessions

Since coordinated HPC and PFC activity during sleep SWRs is implicated in the consolidation of memories (e.g., Peyrache, et al., 2009; Girardeau, et al., 2009; Maingret, et al., 2016; Rothschild, et al., 2017), we compared the incidence of neurons reactivated during post-session sleep (S2) after RC as well as SC sessions. The incidence of neurons modulated by SWR in S2 was significantly greater than S1 in RC sessions (18.4%) than in Training sessions (8.8%; Fig. 7A; χ²=9.25, df=1, p=0.002) and this S2 value was significantly above baseline. The incidence of SWR modulation in S2 was also greater than S1 in sessions with an RC and an SC (Fig. 7A; χ²=7.66, df=1, p=0.006). However, the S2 incidence was not significantly above baseline, while in S1 it was significantly lower. No significant increase in SWR modulation was observed during any sleep sessions prior to or after an SC session (Fig. 7A; χ² test, p>.05). A significantly higher proportion of all neurons were reactivated during SWRs during S2 than during S1, both in RC-only sessions (χ² = 5.16, df=1, p = 0.013; Fig. 7A) and in RC+SC sessions (χ² = 9.12, df=1, p=0.0014; Fig. 7A). However, this difference was not significant in SC-only sessions (χ² = 1.41, df=1, p = 0.166) or in Training sessions (χ² = 2.22, df=1, p = 0.088; dashed lines in **Fig. 7A**).

**Figure 7.**
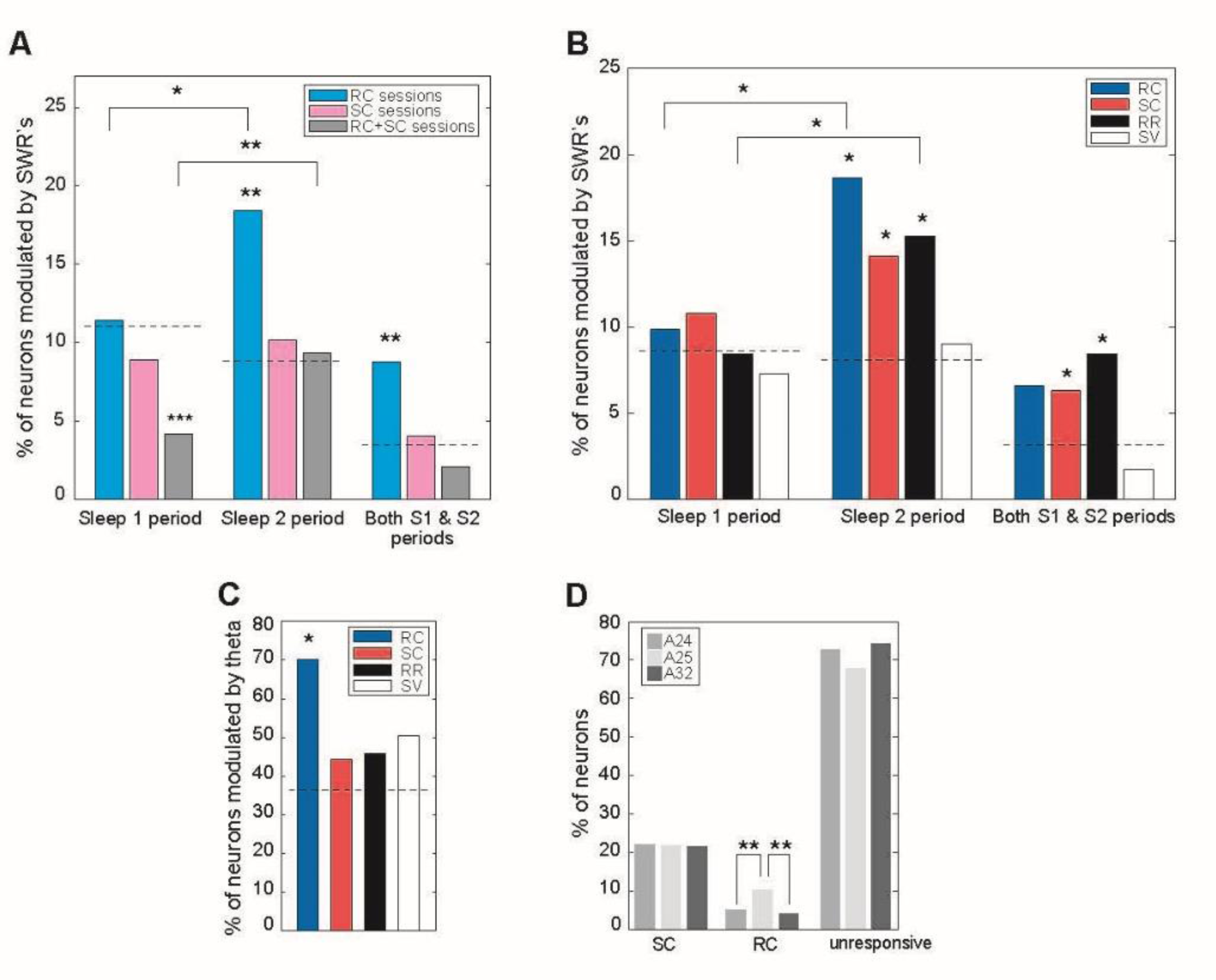
Modulation of ACC activity by hippocampal sharp-wave ripples (SWR) and theta oscillations. **A)** Incidence of neurons with SWR modulation in sleep before and after sessions with a RC, a SC, and both. Dashed lines indicate reference baseline (here, neurons in Training sessions). “Sleep 1 period” includes neurons modulated by SWR in S1 only, as well as in both S1 & S2. Same for “Sleep 2 period”. Stars above single bars indicate a significant difference from relative to baseline (dashed line; unresponsive cells recorded in all sessions). **B)** Incidence of SWR modulation for SC-, RC-, strategy value, and RR responsive cells. Dashed line indicates reference baseline (here, from unresponsive neurons in training sessions). **C)** Incidence of theta modulation according to cell response category. Baseline is proportion of unresponsive neurons in all sessions. **D)** Anatomical distribution of responses. * - p<0.05; ** - p<0.01; *** - p<0.001.

The SWR modulation ratios of reactivated neurons during S1 vs. S2 were indistinguishable for RC sessions (1.41 and 1.39 respectively; rank sum test, p=0.85) and SC sessions (1.60 and 1.41; rank sum test, p=0.085) or RC+SC sessions (1.42 and 1.39; rank sum test, p=0.81). In summary, more ACC neurons were modulated by SWR during S2 than in S1 for RC and RC+SC sessions, but not SC sessions, and there was no significant change in the strength of the modulation.

### Hippocampal SWR modulation of cells with identified responses

The next question concerned whether such activity modulations by hippocampal SWRs were specific to neurons with firing rate changes relative to RC or SC, or to those with other responses, and if, in these specific cases, this modulation was greater in S2 than in S1. Strikingly, a significantly higher proportion of RC-responsive neurons were modulated by ripples during S2 than during S1 (χ2=4.23, df=1, p=0.040; Fig. 7B). This difference also held in RR responsive neurons (χ2=3.85, df=1, p=0.05; Fig. 7B), and was close to significant in SC-responsive neurons (χ2=3.53, df=1, p=0.06). In contrast, this was not significant in SV-responsive neurons (χ2=0.634, df=1, p=0.43). Importantly, none of the groups had a significantly greater proportion of neurons modulated by SWRs during S1 than unresponsive neurons (χ2 test, p>.05). In S2, the incidences of RC, SC and RR neurons were above baseline (χ2=8.4, 7.1 and 4.4, df=1, p=0.004, 0.003 and 0.04). Thus, greater proportions of RC- and RR-responsive cells modulated during SWRs in S2 than S1 were not due to sub-baseline values in S1. Among neurons modulated by SWR during both S1 and S2, SC-and RR-responsive neuron proportions were significantly higher than non-responsive neurons (χ2 = 8.3, df=1, p=0.004; Fig. 7B). This could contribute to maintaining task-relevant information between the two sleep periods.

There were two types of SCs: to rewarded or to unrewarded strategies. The proportions of SWR-modulated neurons in these two groups were not significantly different (S1: 6.7% vs 3.9%; S2: 4.5% vs 8.6%, S1 & S2: 5.6% vs 6.6%, χ2 = 2.99, df=3, p=0.394). Similarly, the incidence of SWR modulation in SV-responsive neurons was not significantly different for rewarded vs non-rewarded strategies (S1: 6.0% vs. 2.6%, S2: 8.0% vs. 2.6%, S1 & S2: 1.2% vs 5.3%; χ2 = 5.20, df=3, p=0.158). As observed in computational models of strategy learning (e.g., Dollé et al., 2018), this is consistent with reinforcing values of correct strategies or weakening values of unrewarded strategies each playing symmetrical roles in ACC-dependent learning, and would thus be equally reactivated during sleep.

A possible confound is that cells with higher firing rates have a greater chance to be reactivated and modulated by SWRs during sleep (Fernando-Ruiz, et al., 2019). Indeed, overall, ripple-modulated neurons here had higher average firing rates (11.9 Hz) during behavior than non-ripple-modulated neurons (5.6 Hz; Kruskal-Wallis test, χ2=155.5, df=1, p=0). However, a two-way ANOVA, with neuron category (SC, RC, SV, RR, non-responsive cells in RC-SC sessions, and non-responsive cells in training sessions) and sleep period as factors, revealed no firing rate difference between neurons modulated by ripples in S1 and in S2 (F=0.13, df=1, p=0.72), and no significant interaction (df=5, p=0.53), despite the differences between S1 and S2 reported above. The neuron category factor was significant (F=3.3, df=5, p=0.006). But the only group with a significantly lower firing rate (RR selective neurons) was the one with a significantly greater incidence of ripple modulation (Tukey-Kramer post-hoc test, p<.05; Supp. Figure 2).

### Modulation of RC and SC responsive neurons by the hippocampal theta rhythm

During all sessions with a RC and/or a SC, about 39% (838/2127) of the ACC cells analyzed were modulated by the HPC theta rhythm (Rayleigh Z tests: p<.05). Importantly, HPC theta modulated a significantly greater proportion of RC-selective cells (70%) than unresponsive cells from all sessions, while SC-, RR-, and strategy value neurons did not (**Fig. 7C**). However there was no significant difference in the strength of modulation between RC- and SC-responsive cells (RC: 6.0%, SC: 6.5%; Mann-Whitney test, zval 0.096, rank sum=3329, p=0.92). Differences in incidence of theta modulation could not be explained by differences in average firing rates in these groups (Supp. Figure 2; Mann-Whitney test, zval=1.234, rank sum=3692, p=0.22). (Differences in running speed were not related to differences in incidence of theta modulation; see Supplemental Data).

### Anatomical distribution of neurons with activity transitions in relation to RC and SC

The proportion of RC-responsive neurons was higher in A25 than in A24 or A32 (**Figure 7D**; χ²=10.7, df=2, p=0.0024). Furthermore, a significantly lower proportion of RC-responsive neurons (7.7%) was located in the dorsal third of A24 (near A32), compared to SC-responsive neurons (21.3%) and unresponsive neurons (19.6%; χ²=7.73, df=2, p=0.0105). The ratio of pyramidal cells to interneurons did not vary significantly among neurons with the respective response types (74% pyramidal cells for RC-responsive, 80% for SC-responsive, 79% for unresponsive; χ²=2.47, df=2, p=0.15). Nor did the concentration in superficial vs deep ACC layers vary for RC or SC (38% were in deep layers).

In summary, more ACC neurons were modulated by HPC SWRs during S2 in RC sessions than baseline, and the incidence in S2 was greater than S1 in these sessions too. Furthermore, modulation by HPC SWRs preferentially occurred in neurons selective for RC, SC and RR, with greater incidence in S2 than S1 for RC and RR. RC-responsive neurons also had a greater incidence of theta modulation than other response types, although all groups had high values. Overall, these results show that modulation by hippocampal theta and SWR was preferentially associated with neurons responsive to changes in task contingencies.

## Discussion

Here, populations and substantial proportions of individual ACC neurons responded rapidly after changes in reward contingencies (RC), or in a fewer trials prior to a SC, and were modulated by SWRs or hippocampal theta rhythms. The experimental design permitted characterization and comparison of the neurons with these two response types in the same data set. In effect, the difficulty of the task permitted comparisons with activity in sessions with no RCs, when the animals spontaneously shifted between different strategies in search for the correct one. Thus, delays between RC and SC were often sufficiently long to permit analyses to distinguish activity changes relative to these respective events, reducing possible confounds. In comparable studies that addressed other questions, RC and SC responses could not always be clearly distinguished (Rich and Shapiro, 2009; Durstewitz, et al., 2010; Karlsson, et al., 2012; Powell and Redish, 2016; Malagon-Vina, et al. 2018; Trouche et al, 2018). Here, we extend upon these and other results by showing that ACC activity also makes abrupt transitions for extradimensional shifts, here between spatial vs. sensory discrimination rules and strategies. Trial-by-trial analyses permitted greater precision than trial block analyses. Thus, the population analyses typical of many studies were complemented here by analyses of individual neurons permitting detection of hippocampal theta and SWR modulation.

### Theta and SWR modulation

Strikingly, a dramatically higher proportion of RC-responsive cells (than SC-, RR-, strategy value-responsive, or unresponsive cells) were modulated by hippocampal theta. This is consistent with a theoretical framework wherein the hippocampal theta rhythm coordinates coherent activity in HPC and ACC for registering experiences. This would lead to synaptic modifications underlying labile memory traces through hippocampal neurons firing sequentially in successive phases of theta cycles (Skaggs et al., 1996; Wagatsuma and Yamaguchi, 2004), and associated PFC activity. SWR events correspond to the replay of sequences of hippocampal activity from the preceding behavioral session, and this modulation could correspond to consolidation of memory traces via transmission to the cerebral cortex, and ACC in particular (Buzsáki, 1989; review: Girardeau and Zugaro, 2011).

Experimental evidence for this is provided by increased reactivation of neurons during SWR’s in S2 compared to S1. During S1, the incidence of hippocampal SWR modulation was indistinguishable among cell groups as well as from baseline values. Furthermore, a significantly higher proportion of in RC- and RR-responsive cells were modulated by SWRs in S2 than S1. This is consistent with a hippocampal contribution to ACC responses to changes in task contingencies, since the hippocampus responds to mismatches between expectations and observations, and novelty (Gray, 1982; O’Keefe, 1976; Vinogradova, 2001; Kumaran and Maguire, 2006; Kafkas and Montaldi, 2018). The proportion of SWR-modulated SC cells was greater than baseline in S2, but was not significantly greater than S1. The greater incidence of hippocampal SWR modulation of RR-responsive neurons is consistent with the enhanced replay of rewarded trajectories in post-learning sleep (Michon, et al., 2019). Interestingly, in an elevated plus maze anxiety paradigm, Adikhari et al (2011) also found increased ventral hippocampal theta modulation of ACC neurons with task-related responses.

In summary, both theta and SWR modulation could reflect selective transmission of hippocampal system signals that would contribute to these ACC neuronal responses. The high incidence of RC- and RR-responsive neurons suggests a role for HPC-ACC coordination in signaling new environmental contingencies. In SC-responsive neurons, hippocampal signals could contribute to elaborating appropriately adaptive behavioral responses.

Those responsive cells not modulated by sleep SWRs might be expected to be modulated by awake SWRs (Tang, et al., 2017). Unfortunately, we did not make reliable recordings of awake SWRs, and thus we could not test this. Furthermore, since modulation was only investigated in neurons with behavioral correlates during mobility, the present work would not have detected cells active during awake immobility when they would be modulated by SWRs (Yu, et al., 2017).

### ACC neuronal response correlates

The RC-and SC-responsive neurons, as well as the RR and strategy value responsive neurons, could participate in ACC’s role in promoting flexible behavior (Granon and Poucet, 2000; Cardinal et al., 2002; Gisquet-Verrier and Delatour, 2006). ACC lesions impair behavioral flexibility in response to a change in the task rule (de Bruin et al., 1994; Birrell and Brown, 2000; Colacicco et al., 2002; Salazar et al., 2004; Lapiz and Morilak, 2006; review: Hamilton and Brigman, 2015). Moreover, ACC damage particularly impairs performance of tasks requiring shifting from one strategy to another, whether the initial strategy has been learned (Granon and Poucet, 1995; Ragozzino et al., 1999a, b) or is spontaneously used by the animal (Granon et al., 1994). Interestingly, here RC-responsive cells were significantly more concentrated in A25 (formerly, IL) than SC-responsive cells and other unresponsive cells. This is consistent with lesion studies showing that A25 lesions impair visual cue reversal learning, while sparing initial acquisition of visual cue discriminations (Chudasama and Robbins, 2003; Li and Shao, 1998). For goal-directed behavior (as well as conditioned fear extinction), A25 has been considered to exert a “stop signal”, and also to suppress representations of action-outcome contingencies from influencing behavior (Barker, et al., 2014). Indeed, an optimal response to a rule change would require cessation of adherence to the previous rule, consistent with the increased incidence of A25 RC responses here.

Powell & Redish (2016) showed that ACC activity transitions were more related to detecting a task rule change, and initiating new behavioral strategies than to initiation of the animal’s behavioral response per se. Here, we extend these results by showing that the communication between HPC and ACC via theta and SWRs is even greater for the detection of task rule changes than for the initiation of new behavioral strategies. Hippocampal contextual information sent to the ACC (and their mutual output structure, the ventral striatum) would be integrated with reward history information to detect task rule changes, perhaps in cooperation with the RR-responsive neurons, as well as A24 (Hyman et al., 2017). The rule shifts would be experienced as the failure to continue to consistently receive reward for a previously successful strategy. This is consistent with the concept of the HPC as a detector of mismatch and novelty, as evoked above. Note that the RC-responsive activity occurred on parts of the maze visited prior to the reward site, thus corresponding to contextual rather than simple post-reward responses. Furthermore, RC responses occurred in a population of neurons with no significant activity correlates with reward rate.

Here, RC and SC associated activity changes in individual neurons were restricted to limited portions of the maze. It is possible that such effects could be masked in population analyses, which average firing over all cells recorded over entire trials. Overall, the population analyses of all neurons showed shifts occurring about 2.5 trials after RC’s and about 1 trial before SC’s. In contrast, the analyses of individual neurons with behavioral correlates modulated by RC or SC showed averaged responses at about 4 trials after the RC and about 3 trials before the SC’s. Since the sampled data sets, calculations and averaging methods are quite different, it is difficult to directly compare the results of the two approaches which suggest that, overall, the population activity responds more rapidly to RC, but that individual neuronal activity “predicts” SCs earlier. Nonetheless, they agree that PFC neurons do anticipate strategy shifts and respond rapidly to rule changes.

In order to learn and perform goal directed choice tasks, one must learn from recent experience concerning the presence or absence of reward and the association of cues and actions. The durations of the behaviorally correlated ACC activity observed here are suited for this since they could be rather brief, or could extend over the order of seconds (cf., Fig. 5). Such activity could serve as buffers for working memory and data processing – not only for cues but also for movements, behavioral sequencing and cue-choice contingencies (Procyk & Goldman-Rakic, 2006; Wang & Hayden, 2017). The ACC could be involved in learning associations of concurrent and sequential events including actions (and not just simple cue-action associations) (Del Arco et al., 2017). These processes could lead to development of reward-based strategy learning. The signals recorded here could hence be considered as instrumental for cerebral processing of the stimulus-action-outcome relations that rules represent.

The RC and SC responses could be informed by modality-specific neocortical inputs relevant to the respective strategies (e.g., visual vs spatial cues for the light-dark vs left-right tasks respectively), as well as by thalamic inputs through cortico-striatal loops (Alexander and Crutcher, 1990; Haber, 2003). The changes in activity levels and formation of assemblies of co-activated ACC neurons can be considered to correspond to functionally active modules that would synchronize with, and activate subsets of downstream basal ganglia neurons to participate in strategy and behavioral choices. Such selection among different rules could also be viewed in terms of neural, attentional and behavioral rechanneling. Indeed, other electrophysiological studies have also reported rodent ACC activity to reflect crucial parameters underlying flexible goal-directed behaviors, such as task-related locomotion (Poucet, 1997; Jung et al., 1998; Fujisawa et al., 2008), reward (Pratt and Mizumori, 2001; Miyazaki et al., 2004), working memory (Baeg et al., 2003), and action-outcome contingencies (Mulder et al., 2003; Kargo et al., 2007). Importantly, the RC-responsive and SC-responsive subpopulations in the ACC are relevant to computational studies implicating rat ACC in action selection (e.g., Hasselmo, 2005; Martinet et al., 2011), and in detecting environmental changes (Caluwaerts et al., 2012). Indeed, our results suggest that the rat ACC is well-positioned to combine contingency change detection and strategy selection functions (Holroyd and McClure, 2015), integrating them during behavior to help build lasting memory traces in conjunction with hippocampal SWRs during sleep.

## Materials and Methods

### Rats

Five Long-Evans male adult rats (225 to 275 g; from the Centre d’Elevage René Janvier, Le Genest-St-Isle, France; RRID: RGD 18337282) were maintained in clear plastic cages bedded with wood shavings. A 12 h/12 h light/dark cycle was applied and all manipulations took place during the light part of the cycle. The rats were housed in pairs while habituating to the animal facility. They were weighed and handled each workday. Prior to pre-training they were placed in separate cages and 14 g of rat chow were provided daily after the pre-training or recording session in order to maintain body weight at not less than 85% of normal values (as calculated for animals of the same age provided *ad libitum* food and water). The rats were examined daily for their state of health and were fed to satiation at the end of each work week. This level of food deprivation was necessary to motivate performance in the behavioral tasks, and the rats showed neither obvious signs of distress (e.g., excessive or insufficient grooming, hyper- or hypo-activity) nor health problems. The rats were kept in an approved (City of Paris Veterinary Services) animal care facility in accordance with institutional (CNRS Operational Committee for Ethics in the life sciences), national (French Ministère de l’Agriculture, de la Pêche et de l’Alimentation No. 7186) and international (US National Institutes of Health, Helsinki Declaration) guidelines.

### Surgery

From the start of the pre-training, rats were placed on the mild food deprivation regime. For at least a week before surgery, rats were habituated to running on the Y-maze by allowing them to forage there for 5-25 minutes daily, with reward available at the end of each arm. Once habituated to the experimental environment (after 7 to 10 days), rats were anesthetized with intramuscular xylazine (Rompun, 0.1 ml) and intraperitoneal pentobarbital (35 mg per kg of body weight). Xylocaine solution was injected under the scalp. A drive containing seven tetrodes (six for recording, plus one as reference) was implanted on the skull above the right medial PFC (anterior-posterior, 3.5–5 mm; medial-lateral, 0.5–1.5 mm). Each tetrode was placed in a 30-gauge stainless steel tube, and the tubes were assembled together in two adjacent rows. Tetrodes were twisted bundles of polyimide coated nichrome wire (13 µm in diameter, Kanthal, Palm Coast, FL). Microdrives allowed independent adjustment of tetrode depth. After retraction of the dura, the rows of cannulae were implanted parallel to the sagittal sinus so that they targeted the superficial and deep layers of the medial bank of the cortex. A separate microdrive containing three tetrodes targeted the ventral HPC (anterior-posterior, -5.0 mm; medial-lateral, 5.0 mm) since the principal projections to ACC originate there (Thierry, et al., 2000). A screw implanted on the occipital bone above the cerebellum served as the reference electrode. The hippocampal tetrodes were lowered to the CA1 pyramidal layer; the depth was adjusted with the help of LFP signs (flat sharp waves, strong ripple oscillations). After surgery, rats recovered for at least 2 weeks while the tetrodes were lowered to reach A24, A32, and A25, as well as the ventral hippocampal CA1 pyramidal layer. Between sessions, tetrodes were gradually lowered to probe different dorso-ventral levels in the ACC.

### Behavioral task

The Y-maze (**Fig. 1A**) was formed by three arms (85 cm long, 8 cm wide with 2 cm high borders) separated by 120 degrees, with a cylindrical barrier at the center. The maze area was surrounded by a cylindrical curtain with no explicit cues on it. The rats were habituated to the maze. The rats had to learn by trial-and-error to adapt their behavior to the current rule in order to maximize the amount of reward received for a given amount of effort. Rats started all trials from the *departure* arm, and after the central barrier was lowered, a light went on at the end of one of the two goal arms selected in a pseudo-random sequence. The rats then had to select one of the two choice arms and then go to the end for reward. Chocolate milk rewards (30 µl) were delivered for choices adhering to the current rule. A photodetector signaled arrivals at the reward trough and triggered a solenoid to release reward when appropriate. On each trial, the reward was available on only one arm. Initially, the animal had to learn that the reward was always located on the right arm, no matter which arm was lit (*Right rule*). Once the animal entered a reward arm the entrance was blocked and it was not unblocked until the rat had visited the reward trough (rewarded or not). The rats had been trained to return to the departure arm after the outcome of the trial (rewarded or not), and the barrier was raised until the next trial. To avoid extensive overtraining, a different rule was applied (Rule Change: RC) after the rat had clearly acquired the current rule (i.e., performance reached a criterion level of 10 consecutive rewarded trials), or 11 rewards in the previous 12 trials. This stringent criterion was applied because some rats tended to engage other strategies after initial acquisition and this assured that the performance had become stable. The rule change was not explicitly signaled to the rat in any way, and could only be detected by the pattern of unrewarded trials. After acquiring the Right rule, the rat then had to learn to go to the lit arm, regardless of whether it was on the left or on the right (*Light rule).* After that was learned, the rats were successively challenged with a *Left rule* and a *Dark rule* (where the unlit arm was rewarded), and then, in the only rat that succeeded in reaching this level (rat 20), the sequence of task rules was started again from the beginning (*Right*, *Light*, etc.). All rule changes thus qualified as extra-dimensional, engaging spatial or visual cues respectively. No intra-dimensional shifts (reversals) were imposed such as *Right rule* -> *Left rule*, or *Dark rule* -> *Light rule*, or their inverses, since these have been reported to not require medial PFC function (Birrell and Brown, 2000).

Rats were trained in daily sessions. Each session consisted of 10 to 60 consecutive trials, stopping when the rat ceased performing. Each session began with the same rule as the final trials of the previous session. Since several sessions were sometimes required to learn certain task rules, there were sessions where no shift in the task rule was imposed. Thus, we will distinguish *shift sessions* (i.e., sessions where a shift in the task rule occurred) from *non-shift sessions*.

### Data acquisition

Spike waveforms were filtered between 600 and 6000 Hz, digitized with a Power1401 interface (Cambridge Electronic Design, UK) and time-stamped. For this, 32 samples at 32 kHz (1 ms total) were recorded whenever the signal exceeded a manually set threshold (8 pre-trigger and 24 post-trigger samples). The signal recorded from the same tetrodes was also passed into a low-band filter (between 0.1 and 475 Hz) in order to extract LFPs. The timestamps of behavioral events (e.g., photodetector crossings) were integrated with the spike data online. A video camera was synchronized with the data acquisition software and monitored the consecutive positions of the animal during the experiment. The video tracking data of the instantaneous position of the animal were acquired using MaxTRAQ^®^ (Innovision Systems, Columbiaville, MI) software.

For action potential spike discrimination, the data were processed with a custom Python script for Principal Component Analysis. Then, the Klusters and Klustakwik software (Harris et al., 2000; Hazan et al, 2006; Buzsáki Lab, RRID:SCR_008020) carried out the spike-sorting using the expectation-maximization (EM) algorithm (Celeux and Govaert, 1992). The classifications were then revised manually. Parameters of the EM algorithm were intentionally chosen to extract a high number of clusters (typically from 15 to 60 from the multiple tetrode array). Then those clusters likely to belong to the same unit were merged in manual online analyses.

At the end of experiments, a small electrolytic lesion was made with cathodal current through one wire of each tetrode (25 µA for 10 s). Rats were euthanized with sodium pentobarbital and perfused with saline then formaldehyde solution (4% v/v). Lesion sites were detected in Nissl stained 60 µm sections. Positions of recorded neurons were determined with respect to the lesion site by interpolating the measured descent of the electrodes. Electrode sites were reconstructed in three dimensions with the Neurolucida^®^ system.

### Identification of putative pyramidal cells and interneurons

Putative interneurons and pyramidal cells in ACC were distinguished according to spike width with the method from Barthó et al. (2004). The distribution of spike widths for all ACC neurons was recorded and was bimodal (see Supplementary Figure 6 in Benchenane et al, 2010). Cells with spike widths inferior to 0.3 ms were classified as putative interneurons, whereas those superior to 0.35 ms were considered as putative pyramidal cells.

### Behavioral analyses

Each trial was characterized by parameters including the current task rule (right, light, left or dark), the position of the light (right or left), the arm chosen by the animal (right or left), and the outcome of the trial (rewarded or not).

Two principal events were the bases for the analyses of behavioral and neurophysiological data from each trial: *Start* and *Arrival*. The trial Start time was extracted from the video tracked position data and was defined as the onset of initial forward acceleration in the departure arm. The Arrival event was defined as the instant when the animal first blocked the photodetector at the reward reservoir of the selected arm.

Next, to systematically categorize the animal’s behavior in terms of the “strategy” it followed, the behavioral data from all of the sessions were concatenated for each animal. A custom script scanned these files and extracted series of trials where the animal’s behavioral strategy was consistently stable (“blocks of trials”). The strategies considered in the behavioral analyses included: 1) Going to the right arm regardless of the visual cue (“Right” strategy); 2) “Left”; 3) Going to the lit arm, regardless of whether it was to the right or left (“Light” strategy); 4) “Dark”; and 5) Spatial “Alternation” (not rewarded) (see Figure 1A). Trials not included in such blocks are considered as a sixth “Undefined” strategy, while the latter five are termed “Defined”. We systematically searched for higher order strategies (e.g., left-left-right-right-left-left, etc.) but found insufficient evidence for them. Note also that a single trial could be compliant with two strategies (for instance Left and Dark) or even three strategies (for instance Left, Dark and Alternation), but the randomization of the light positions assured that blocks of 4 or more trials could not comply with more than one strategy.

To determine the minimum number of strictly compliant trials to be considered a strategy block, we performed simulations consisting of 1.6x10^7^ random runs. Each of these runs included the 3322 trials of the data set, with the same sequence of rules and lit arm positions, but the arm choices were randomly generated. These simulations showed that for blocks of length 6 or greater the only 5.4% of random arm choices would belong to such a block for each single strategy. Thus, 95% of random moves would not belong to such a block. Hence, we considered a block to be strictly compliant to one of the five Defined strategies if it contained a continuous sequence of at least 6 trials such that all the rat’s choices were compliant with that strategy. This is the same, or even more stringent criterion than used in this type of study (e.g., Kaefer, et al., 2020; Lapiz-Bluhm, et al., 2009, Tait, et al., 2017).

Since rats occasionally made non-compliant choices within long runs of compliant trials, we also considered as strategy blocks those sequences containing at least 9 trials with only one non-compliant (NC) trial. For example, if the animal’s choices were compliant (C) with a strategy for 6 trials, NC for 1 trial, and then C again for 2 trials, the whole block of 9 trials is considered as a “leniently compliant” strategy block. Following the same principle, we included blocks of CCCCC-NC-CCC (and its inverse), or CCCC-NC-CCCC. Longer sequences might include several NC trials providing that these rules were respected. In simulations as described above, the probability that a random choice would belong to a leniently compliant block for each of the strategies was only 8.6%; 91.4% of random moves would not belong to such a block.

To gauge the risk that sequences detected with these criteria could have occurred stochastically, the incidence of leniently compliant sequences was computed in simulations of 1.6x107 random runs. Each random run included the 3322 trials of the data set, with the same sequence of rules and lit arm positions, but the arm choices were randomly generated. These simulations yielded values of the mean, median and 95% confidence intervals for the percentages of trials that would have stochastically appeared in leniently compliant blocks of length greater than or equal to 6 trials (Table 1). For all of the strategies, the incidence of experimental trials in blocks compliant to the respective strategies are outside of the confidence interval limits, inconsistent with fortuitous compliance by means of stochastic choices. Similarly, the incidence of trials following any defined strategy (78.2%) greatly exceeded the confidence interval upper bound (43.7%).

**Table 1.** Incidences of trials in leniently compliant blocks of length >5. At left are results for the simulations described in the text. Actual values are from animal data. Note that blocks compliant with different specific strategies could partially overlap, so that the sum of trials for the five strategies do not equal that of Defined trials.

| Simulation results (% trials in blocks) |  |  |  |  | Actual values |  |
| --- | --- | --- | --- | --- | --- | --- |
| Strategy | Mean | Median | Lower bound (2.5%) | Upper bound (97.5%) | % | n |
| Left | 8.6 | 8.6 | 5.8 | 11.6 | 27.0 | 896 |
| Right | 8.6 | 8.6 | 5.8 | 11.6 | 22.0 | 730 |
| Light | 8.6 | 8.6 | 5.8 | 11.6 | 23.4 | 776 |
| Dark | 8.6 | 8.6 | 5.8 | 11.6 | 6.9 | 229 |
| Alternation | 8.6 | 8.6 | 5.8 | 11.6 | 13.8 | 459 |
| Defined | 38.9 | 38.9 | 34.2 | 43.7 | 78.2 | 2599 |
| Undefined | 61.1 | 61.1 | 56.3 | 65.8 | 21.8 | 723 |

To verify that the results of these analyses were not dominated by very long sequences that were clearly not stochastic, we further examined smaller blocks of 11 trials or less (which composed 99.1% of the trials in randomly generated choices). Over these respective block lengths, 40% of the actual values fell outside the p<.05 confidence limits, again providing evidence against stochastic choices.

The performance of the animals prior to initial rule acquisition was also inconsistent with stochastic behavior. Four of the five animals reached criterion for the R rule in the very first training session (cf., values in Results “Behavior”), corresponding to fewer trials than expected by chance. During the initial acquisitions, the incidence of occurrence of sequences compliant with a rule currently not being rewarded also extended beyond the lower or upper bounds of the confidence intervals of the randomized data simulation.

*Strategy changes (SC)* are defined as two consecutive blocks where the rat followed different strategies (e.g., the animal performs a series of 10 trials following the *Light* strategy, always going to the lit arm, and then performs a series of 6 trials following the *Alternation* strategy). We define *extradimensional shifts* as SC between spatial and visual strategies (or vice versa), such as Left then Dark strategy.

### Experimental design and statistical analyses

Neuronal activity was recorded as the rats performed in a completely automated maze, or during sleep immediately preceding and following this. Each neuron served as its own control and activity levels were compared before vs. after events with analyses and statistical tests as described in the following sections. The tests include bootstrap Monte Carlo analyses, ANOVA, t-tests, Kruskal-Wallis, Mann-Whitney, χ2, and Rayleigh Z tests. The specific applications of these tests are explained below and in the Results section. (Insufficient sampling precluded analyses of REM sleep).

### Activity transition analyses at the population level

An unbiased automatic decomposition of the trials in each session into blocks determined whether the onsets of transitions in neural activity are most closely temporally associated to behavioral (SC) or task (RC) changes. Similar to Powell & Redish (2016), for each trial in a given session, we computed a population vector from the firing rate of each given neuron averaged over the entire trial (from *Start* until *Arrival*) from all PFC cells recorded simultaneously. Each population vector was z-scored in order to give the same weight to cells with low and high firing rates. We then computed the correlation matrix where each element *E_ij_* is the correlation between population vectors of trials *i* and *j* for all trials of the session.

When displayed graphically, blocks of high values in the correlation matrix indicate series of trials with correlated population activity. We performed an automated search for such blocks by detecting the best decomposition of the matrix into blocks, i.e., finding the optimal partition of blocks of trials along the diagonal of the matrix, so that eliminating elements outside the blocks minimizes the amount of information lost. Information loss was measured as the percentage by which the sum of absolute values of the elements in the matrix is reduced after exclusion of the elements outside the blocks. To prevent the trivial decomposition (i.e., a single block encompassing everything) from being systematically considered as the one that minimizes the loss (since it has no loss of information), we rejected it if at least one decomposition into 2 or more blocks led to a loss of less than 1/3 of the information (i.e., if the information present within the blocks was superior or equal to 2/3 of total information within the matrix). Otherwise, no decomposition was retained, which was interpreted as no substantial change in population activity having occurred during that session.

In order to find the best decomposition into two or more blocks, we tested all possible decompositions of each matrix into blocks (with each block including at least 3 trials and at most 2/3 of the trials of the session). Then for each decomposition, we projected the original matrix onto the considered blocks (which are concentrated around the diagonal of the matrix), hence setting to zero the elements outside the blocks. This tests the hypothesis that the correlations of activity between blocks are negligible, as if the population activity has sharp state changes between blocks of trials. Then, for each decomposition, we measured how much information was lost, and retained the one which minimized this amount.

We developed this approach to decompose the population cross-correlation matrix rather than use k-means clustering employed by Powell and Redish (2016) because it was not clear that the data satisfied the requirements of k-means. Firstly, k-means is based on the principle of spherical clusters that can be separated in such a manner that the means approach the centers of the respective clusters. Furthermore, the clusters should have comparable sizes. But, here, we expected sharp transitions between series of trials with respectively homogeneous firing rates, and the transitions could occur early or late in the session, creating groups of unequal size. Furthermore, it was not clear how many clusters to preset in the k-means algorithm since RC and SC could each occur one or more times in the same session, and the recorded populations only had limited numbers of neurons, leading some of these to not respond to some RCs or SCs. There was also concern that the algorithm would stop at local minima, rather than at more optimal solutions.

### Estimating timing of population activity transitions relative to RC or SC events

In order to quantify the possible relationship between an activity transition detected on a particular trial *i* with a RC or SC event occurring on trial *j*, the difference between *i* and *j* was taken if there was only one such event. If there were two or more events of the same or different types (i.e., RC or SC), the one occurring the fewer number of trials from the activity transition was selected. Possible ambiguities could occur when a RC was rapidly followed by a SC. As a control, analyses were first performed only on sessions where only one such event occurred. Activity changes associated with RC were only considered for the first trial after a conflict between the reward contingency and the previous rule was detectable (since individual behavioral choices can be consistent with more than one rule).

### Event-locked analyses for activity transitions

Activity in time windows bracketing salient trial events were compared in trials prior to and after RC and SC. The four periods included: *pre-start*, in the time window [*Start* – 2.5 sec; *Start*]; *post-start* [*Start*; *Start* + 2.5 sec]; *pre-arrival* [*Arrival* – 2.5 sec; *Arrival*]; and *post-arrival* [*Arrival*; *Arrival* + 1.25 sec]. The post-arrival period was restricted to 1.25 s in order to restrict to the period when the rats were immobile irrespective of whether the trial was rewarded or not. This avoided possible confounds with different motor behaviors in rewarded and non-rewarded trials. In 96.4% of all trials recorded in the five animals (including rewarded and unrewarded trials), the rats remained at the reward site for at least 1.25 s.

The Wilcoxon-Mann-Whitney test examined differences in neuronal firing rate around RC and SC events. This was tested for each of the four trial event windows (with Bonferroni corrections for multiple comparisons).

To confirm and expand upon these analyses, a bootstrap Monte Carlo analysis was also performed for each cell, again with separate tests for each of the four periods. In all cases, the firing rates were compared between trial blocks before and after the RC or SC. Then the activity of these trials was randomly re-attributed to two surrogate sets with the same numbers of trials, and the difference was taken. This shuffling procedure was repeated 5000 times and yielded a distribution from which only those differences in firing rate with p-values falling within the upper or lower 2.5% of the shuffled distributions were considered as significant transitions in neural activity. To quantify the incidence of spontaneous changes of firing rate of PFC neurons, in sessions where the rat consistently performed a single strategy, the first half of the session was compared to the second half with the same bootstrap Monte Carlo analysis. Since the latter tests (reported in Results) showed that a non-negligible number of significant transitions in firing rate occurred in the sessions with no strategy or rule changes (or other apparent behavioral differences), further complementary analyses were performed.

### Estimating timing of individual neuron activity transitions relative to RC or SC events

The preceding population analyses group data from different maze positions. It presumes that activity shifts will only take place at RCs or SCs. Another analysis without this *a priori* assumption examined each neuron’s activity to determine on which trial (if any) there was the maximal significant difference between firing rates in the block of trials before and after that trial. The nonparametric analysis developed by Fujisawa et al. (2008) constructs bootstrap confidence intervals of positional spike density functions. The null hypothesis posits no difference in firing between trials before and after the event of interest (here, RC and SC). Sessions were divided into all possible pairs of sets of contiguous trials, with a minimum of four trials in each set. Thus, if a session had 50 trials, trials 1 to 4 were compared with 5 to 50, 1 to 5 vs 6 to 50, 1 to 6 vs 7 to 50, […], through trials 1 to 46 vs 47 to 50. Spike density functions were determined for each of the pairs of sets of trials.

Here data could only be analyzed for positions that the rat occupied on every trial of the session, and other data (particularly on the start arm, where the rats did not always go to the end) were excluded. Data were divided into spatial bins of length 1.75 cm along the linearized trajectory from starting point to reward site (each corresponding to 2.5 pixels in the video image; x-axis of 1^st^ and 2^nd^ columns of **Fig. 5**).

For each pair of trial blocks, a bootstrap Monte Carlo analysis randomly redistributed all of the trials in the session to surrogate pairs of sets with the same respective numbers of trials as the original sets. For each surrogate set a spike rate function across the spatial bins was taken as a set of points on the maze where the spikes occurred and this was transformed to a spike count function with a Gaussian kernel whose bandwidth is 2 bins (= 3.5 cm). Dividing the spike count functions by the time spent in each position over all of the trials for the respective parts of the session yielded the spike rate functions. The pre-versus post-transition difference was then computed for each block of trials.

To determine the statistical significance of these rate differences, the distribution of the rate difference statistics was estimated by randomly reassigning the data from each trial into two surrogate groups 5000 times. For each of the spatial bins, a pointwise confidence band with a p-value of 0.05 was assigned to the 125 greatest positive and negative differences respectively (a two-way test). Since the pointwise confidence band is computed at multiple points, the confidence level must be corrected for these multiple comparisons. This is achieved by computing the global confidence band: first, we calculate the “global confidence level”, taken as the percentage of resampled rate differences whose values are not all inside the area limited by the pointwise band. The procedure is repeated to build the pointwise confidence band by gradually decreasing the value of the pointwise confidence limit until the global limit is equal to 0.05. Then, a maze zone is considered to have significant difference in pre-versus post-transition firing only if this difference crosses both the global and pointwise bands, but the extent of this zone is determined only by the points where it lies beyond the pointwise band (Fujisawa et al, 2008; also see Catanese et al., 2012).

To control for possible spatial selectivity in the ACC responses (Euston et al., 2007), analyses were repeated separately using only data from either left or right choice trials in sessions where sample sizes were large enough (see also Lindsay, et al., 2018). This showed that 43 (73%; or 34% of the total of 127 neurons) retained significant SDF differences when either left or right choice trials were analyzed alone, and thus their transition related activity changes are not confounded with location-selective firing.

### Multiple regression analysis of single-unit activity with strategy value and reward rate

In order to control for potential confounds in putative RC- and SC-responses, we constructed several regressors, and then tested for correlation with single cell activity with a multiple regression analysis.

We first built a simple and parsimonious (without any free parameters) regressor mimicking progressively learned strategy values in computational models (Dollé et al., 2018). This consisted of the estimated value of the three observed strategies: V_Right_, V_Light_ and V_Alternation_ (V_Left_ being equal to -V_Right_, and V_Dark_ = -V_Light_). These values were initialized at 0 at the beginning of each new session. After each trial, the value of a strategy X was increased by 1 if the animal’s behavior was consistent with it and got rewarded, decreased by 1 if it was inconsistent and got rewarded. Furthermore, the value was decreased by 1 if it was consistent but unrewarded, and increased by 1 if it was both inconsistent and unrewarded. For example, if at trial N the animal went to the left arm which was lit, and received a reward, then V_Light_ was increased by 1 while V_Right_ was decreased by 1.

We then built a second regressor representing the reward rate (RR), computed over a 6-trial sliding window. We finally also included regressors representing the lit arm on the current trial (L; 0 for left or for 1 right), arm choice (C; 0 for left or 1 for right), and reward (R; 0 or 1).

The spike rate y(t) in trial t was then analyzed using the following multiple linear regression model:

y(t) = ρ_0_ + ρ_1_V_Right_(t) + ρ_2_V_Light_(t) + ρ_3_V_Alternation_(t) + ρ_4_RR(t) + ρ_5_L(t) + ρ_6_C(t) + ρ_7_R(t)

where ρ_i_ (i ∈ {1..7}) are the regression coefficients.

Following Seo and Lee (2009) and Khamassi et al. (2015), we applied a restrictive significance threshold permitting only 5% significance after permuting the trial order 10,000 times (bootstrap method) so that this regression model would not suffer from violation of independence between regressors.

### LFP analyses

Each of the mid-ventral hippocampal tetrodes was electrically connected in a single-electrode configuration (all channels shorted together) and used for LFP recordings. The screw implanted in the occipital bone above the cerebellum was the reference. Local field potentials were sampled and stored at 2 kHz. Theta was filtered at 5-10 Hz. Theta modulation of cell activity was detected with the Rayleigh test with a criterion of p<.05. Detecting SWRs and measuring SWR modulation of cell activity employed the methods of Peyrache, et al. (2009). For the latter, PETH’s plotted cell activity relative to ripples with 100 ms bins. A t-test compared the central bin to the baseline value.

For information about and access to the recording data base, see https://crcns.org/data-sets/pfc/pfc-6/about-pfc-6

## Supporting information

Supplementary Annex

## Acknowledgments

We thank Prof. J.-M. Deniau, Drs. A.-M. Thierry, Y. Gioanni, M.B. Zugaro, R. Todorova, and E. Cerasti for valuable discussions, S. Doutremer for histology, F. Maloumian for help with figures, N. Quenech’du for help with the 3-D anatomical reconstructions, Y. Dupraz for mechanical engineering and Prof. A. Berthoz for support throughout the project.

## Data availability

The electrophysiological data are available at Adrien Peyrache, Mehdi Khamassi, Karim Benchenane, Sidney I Wiener, Francesco Battaglia (2018); Activity of neurons in rat medial prefrontal cortex during learning and sleep. CRCNS.org http://dx.doi.org/10.6080/K0KH0KH5

## Supplementary Data

In 341 neurons (83% of the 411 selected neurons; 36% of the total pool of 944 responsive neurons) there were SDF differences on all trials taken together and also on either left choice trials alone or right choice trials alone. Among the 331 neurons that could be analyzed in the 18 RC+SC sessions, 140 (42.3%) showed a significant shift in activity (Monte Carlo, p<0.05, two-tailed). Among these, 121 (121/140: 86.4%; or 36.6% of the total of 331 neurons) retained these differences on either left or right choice trials alone. These neural activity transitions in 83 neurons occurred fewer trials from an SC (and thus will be considered “SC-responsive”), while 90 were fewer trials from an RC (thus considered “RC-responsive”), while in 9 neurons this could not be determined unequivocally. Here individual neurons could show two activity transitions in close proximity to a RC and a SC respectively (62/418, or 15%, of SC-responsive neurons and 62/145, or 43%, of RC-responsive neurons). Because of this ambiguity, analyses were further pursued for SC-only and RC-only sessions only.

For each session with a RC or a SC, the significant SDF values of trial-by-trial bootstrap Monte Carlo analyses of the neurons were plotted as distributions across trials for different spatial positions on the maze, as shown in the second column of **Fig. 5**. This generally revealed smooth progressions of values with maximal or minimal values. However, the trial with the peak SDF value (e.g., trial 8 in **Fig. 5A**) did not necessarily represent the most marked intertrial activity transition (e.g., between trials 22 and 23 in **Fig. 5A**), because of non-monotonic trial-by-trial progression of SDF values.

To verify that the latter results are not confounded by other behavioral or task related parameters, we re-examined the beta distributions for RC-responsive and SC-responsive neurons, excluding those that also showed behaviorally correlated activity including left vs right reward arm choice, correct vs error trial, and light vs dark reward arm choice. In RC-only sessions, the beta distribution means of SDF values was 4.7±3.1 trials after the first unrewarded trial providing evidence for the RC. For SC-only sessions the mean was 2.3±1.6 trials before the SC. In RC+SC sessions, the mean was 4.1±2.9 trials after the RC while the mean was 1.8±1.4 trials before the SC. All of these results are consistent with the above analyses.

The difference in the proportion of theta-modulated cells in RC-only vs SC-only sessions was not due to differences in running speed (which affects theta). The mean velocity of animals during RC-only sessions (36.0±3.8cm/s), SC-only sessions (35.9±1.7cm/s), RC+SC sessions (38.4±2.3 cm/s) and Training sessions (31.1±2.7cm/s) were not significantly different (Kruskal-Wallis test, χ²=2.3, df=3, p=0.5125). There also was not a significant difference between RC-only and SC-only sessions in the change in running speed between the 5 trials before and the 5 trials after the RC or SC event (RC-only sessions: -8.0±4.7 cm/s; SC-only sessions: -3.8±1.8 cm/s; Kruskal-Wallis test, χ² = 2.78, df=2, p=0.25). Thus the greater incidence of theta modulation in RC-responsive neurons is not likely to be confounded with differences in speed changes between RC and SC. Furthermore, note that a recent study showed very minor impact of movement parameters on the activity of individual PFC neurons (Lindsay, et al., 2018; see also Powell and Redish, 2016).

## Supplementary Figures

**Supplementary Figure 1.**
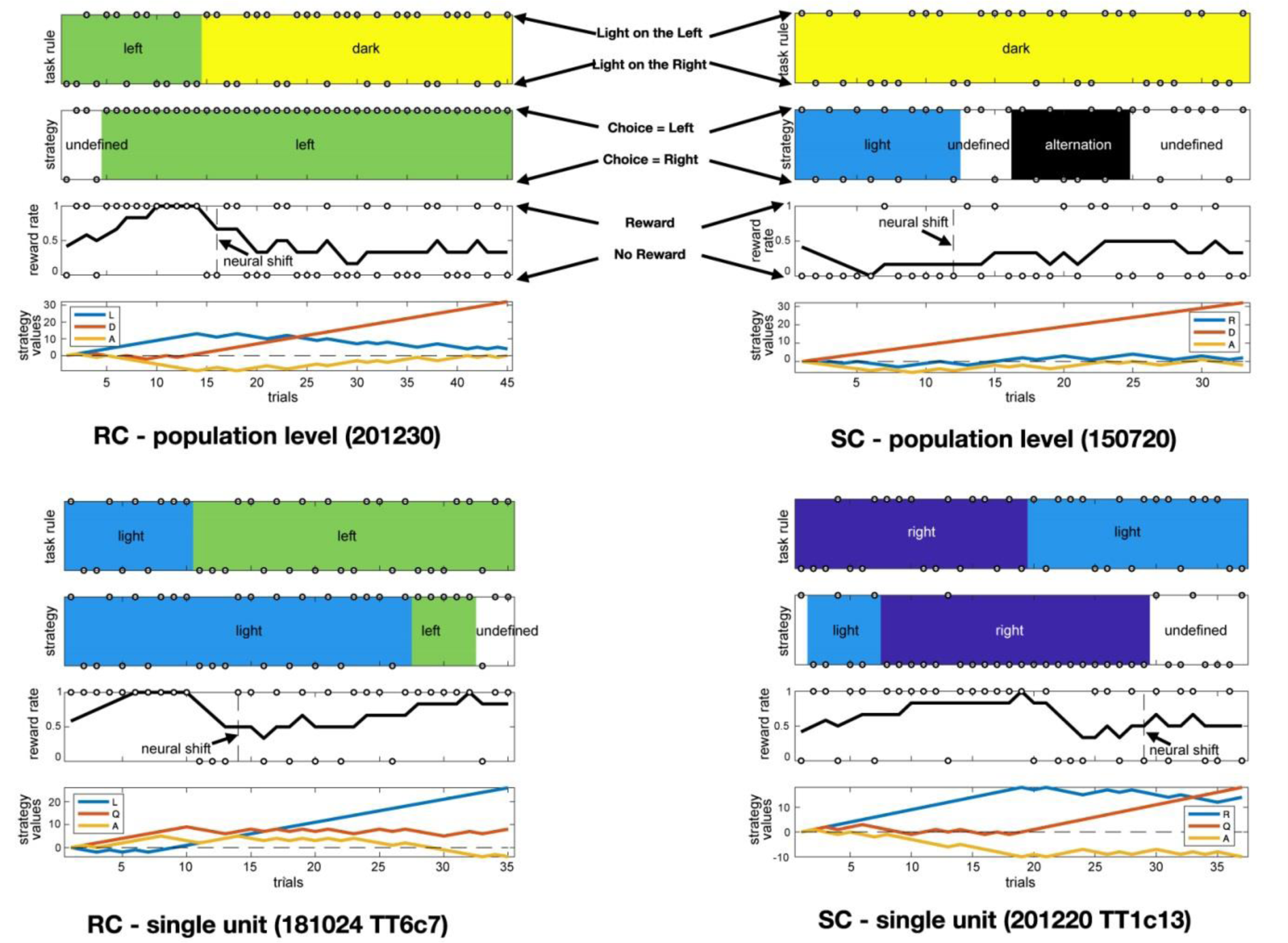
Examples of neural shift events illustrated relative to the rat’s behavior, and other potentially relevant information such as reward rate and strategy values. Top, neural shifts detected at the population level (shifts between blocks within population activity matrix) in two rats, Rat 20 and Rat 15. Bottom, neural shifts in single neurons in two rats: Rat 18 and Rat 20. R=Right strategy, L=Left strategy, Q=Light strategy, D=Dark strategy, A=Alternation strategy.

**Supplementary Figure 2.**
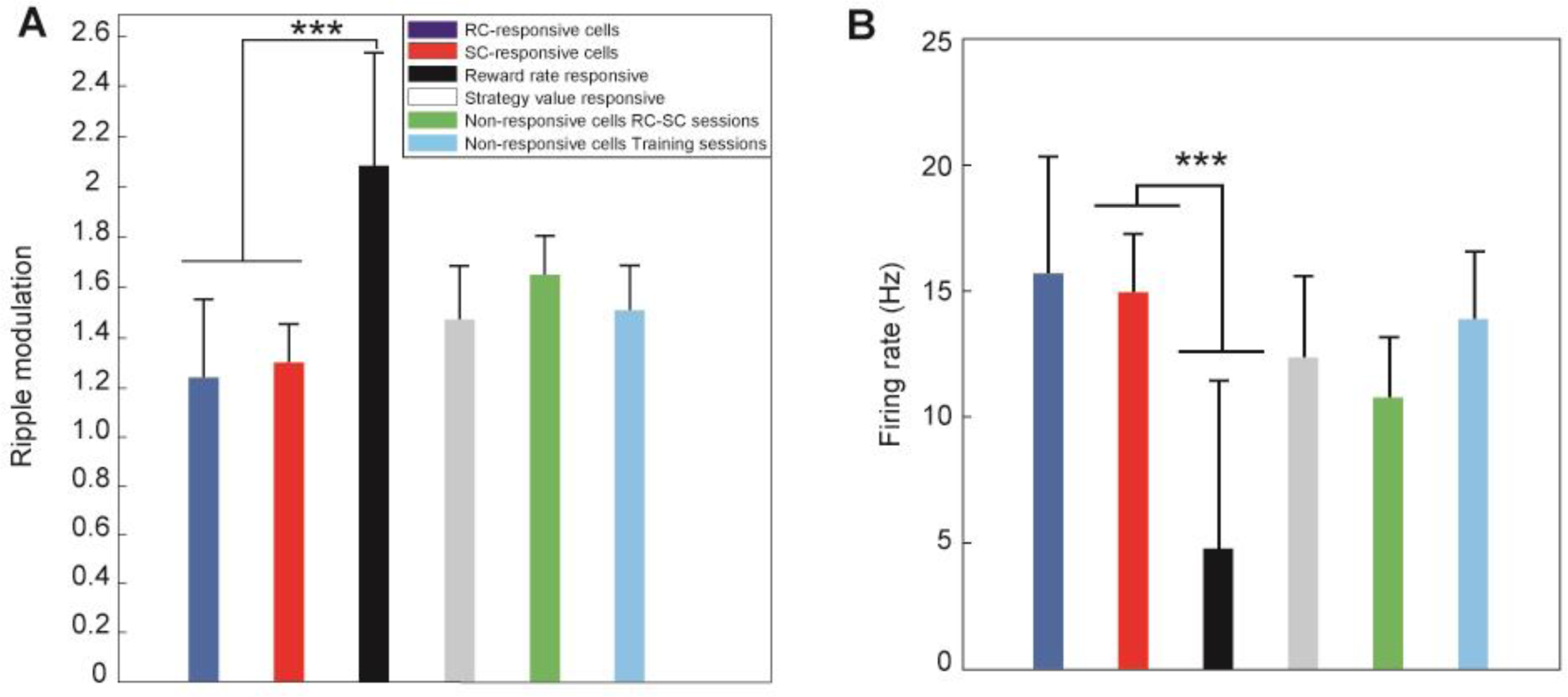
A) Amplitude of ripple modulation among the cell response types before and after sessions. (B) Control for firing rates ***-p<0.001 (Chi-square test)

