## Supplementary Annex for "Rat anterior cingulate neurons responsive to rule or strategy changes are modulated by the hippocampal theta rhythm and sharp-wave ripples"

#### Preliminary statements

The goal of this work is to provide a descriptive analysis of animals behavior in a binary rewarded decision task. Descriptive means that we don't try to explain why or how the animals perform the task, though we have to specify several explicit hypotheses, as follows, which are all of course debatable:

(H1) We assume that each animal decision is based on a finite number of information, which are all available at the decision time (i.e. the behavior is causal )

(H2)We assume that all the cues are binary variables (or can be reformulated as a set of binary variables). As a counterexample of this assumption, the animal decision might be instead based on the accumulated reward obtained from the beginning of the experimental session.

(H3) We suppose that all these cues are also known to the experimenter, and correctly recorded. Note: this hypothesis is quite strong; animal decision might depend on uncontrolled or unrecorded cues. This issue could be eventually handled in the permissive analysis (see below)

(H4) We suppose that at each decision time, the animal will apply a deterministic rule (*i.e.* a Boolean function of the set of binary cues). Again, this is quite a strong hypothesis, which will be relaxed in the permissive analysis.

(H5)We admit that at some trials (the transition trials), the animal switches to a different rule, drawn from a prespecified set of rules. We will not try to explain why the transition occurs, but only when it is likely to occur. That's why the analysis is in some sense only descriptive and not subjective.

(H6) Obviously, the number of transitions has to be restricted in some way. If not, the mere sequence of observed animal decisions would be the best result of the analysis. In the following, we will assume a *constant transition probability* $\varepsilon$ $< 0{:}5$. We will show that, according to the non permissive analysis, maximizing the posterior probability of sequence of rules and transition (given the sequence of observed decisions) is equivalent to minimizing the number of transitions.

#### 1 The non permissive analysis

Let $M^t$ the observed animal decision at trial t with $1 \leq$ t $\leq$T , $R^t$ the unknown rule (or strategy), one element of the set of allowed rules $\mathcal{R}$, applied by the animal at trial t, and $C^t$ the set of binary cues available to the animal (and recorded by the experimenter) at trial t. In the non permissive analysis, we apply a Dirac likelihood distribution, as follows:

$$P(M^t = m \,|\, R^t = r; C^t = c) = m \wedge r(c)$$

This equation means that the likelihood is equal to 1 when the animal decision is exactly predicted by the application of the rule r to the set of cues c, otherwise the likelihood is equal to 0.

According to (H6), the transition probability between rules is specified by:

$$P(R^{t+1} = r' \neq r \,|R^t = r) = \varepsilon/(n-1)$$

$$P(R^{t+1} = r \,|\, R^t = r) = 1 - \varepsilon$$

where n is the number of allowed rules and $\varepsilon$ is the (unique) constant parameter used in this method. We also assume that the prior distribution on R is uniform (no rule is “privileged”), so that:

$$P(R^1 = r) = 1/n$$

The posterior distribution over the whole sequence of rules is given by:

$$P(R^{1:T} \,|\, M^{1:T},\, C^{1:T}) \propto P(M^1 \,|\, R^1,\, C^1) \prod_{t=2:T} P(R^t \,|\, R^{t-1}) P(M^t \,|\, R^t,\, C^t)$$

The problem is to find the sequence of rules which maximizes the posterior probability, given the observed decisions and the recorded cues, $i.e.$:

$$r_{opt}^{1:T} \in Argmax\, P(R^{1:T} \,|\, m^{1:T},\, c^{1:T})$$

Note that several sequences could have the same probability, therefore the analysis might eventually provide several optimal solutions.

To solve this problem, we first remark that if for any single trial the rule is not compatible with the observed decision, $i.e. \exists t : m^t \neq r^t(c^t)$, the whole sequence has a null probability, hence the term “non permissive”. We then constrain the rules to belong to the series of subsets $S^t$ defined as the subsets of compatible rules at trial t:

$$r \in S^t \Leftrightarrow m^t = r(c^t)$$

We also remark that the transition probability at each trial can have only two values: either $\varepsilon/(n-1)$ if there is a rule change, or $1-\varepsilon$ if not. It follows that the Log posterior probability of the whole sequence of rules (LP(R)) depends only on the number $n_T$ of transition trials (providing that all rules belongs to the compatible subsets):

$$LP(R) = LogP(R^{1:T} \,|\, m^{1:T},\, c^{1:T}) = cste\text{-}n_T Log(\frac{(1\text{-}\varepsilon)(n-1)}{\varepsilon})$$

Providing that $\varepsilon < 0.5 \leq \frac{n-1}{n}$, an optimal sequence of rules is therefore a sequence of compatible rules ($i.e.$ $r^t \in S^t$) with a minimal number of transition $n_T$. To compute the minimal $n_T$ and the sequence of optimal rules, we compute the sequence of rule subsets $Q^t$ as follows:

(1) Initialize $n^T = 0$ and $Q^1 = S^1$

(2) For t = 2 to t = T compute $Q^t = Q^{t-1} \cap S^t$ and continue while $Q^t \neq \emptyset$

(3) If $Q^t = \emptyset$, t is a transition trial, then increment $n_T$ and reset $Q^t = S^t$

(4) Between two transition trials $[t_0,\, t_1 - 1]$ (or between the first trial and the first transition $[t_0 = 1,\, t_1 - 1]$; or between the last transition and the last trial), we have $R^t = r$ where r can be freely chosen in the subset $Q^{t_1 - 1}$.

Interestingly the non permissive analysis, though formalized in probabilistic terms, reduced to a logic algorithm, and the resulting optimal solutions do not depend on the parameter $\varepsilon$. However, it clearly depends on the set of allowed rules $\mathcal{R}$ (see results). One has also to keep in mind that the output of this algorithm is one optimal solution; one cannot exclude that there are other optimal solutions, with different transition trials. Suppose for instance that the algorithm output is:

$$aaaaaaaaaabbbbb$$

The rule a is valid for the first 10 trials, then one has to switch to rule b for the remaining 5 trials. Let's suppose that the rule b is not valid for the first 5 trials, but thereafter becomes valid for the remaining 10 trials. It follows that the sequence:

$$aaaaabbbbbbbbbb$$

is as optimal as the first one. It would have been the output of the same algorithm working backward from trial T to trial 1.

#### The Reduced Rule Set Problem

Suppose that we get an optimal solution $r_{opt}^{1:T}$ by taking rules from a set $\mathcal{R}$. The question is: is there a minimal set of rules $\mathcal{R}_{min} \subset \mathcal{R}$ such that, but constrained rules to belong to the reduced set of allowed rules, we obtain exactly the same number of transitions $n_T$ than with a full set of rules ? Unfortunately, this problem is NP complete, and therefore one cannot build the true $\mathcal{R}_{min}$ in a reasonable time. Fortunately, there exists an approximate algorithm (a "greedy" algorithm) which provides a reduced rule set (RRS) which is generally a good approximate of the minimal set.

The principle of the greedy algorithm is the following:

(1) one builds the list of $n_T + 1$ subsets of valid rules with the forward algorithm described before:

$$\{Q^{t_1-1}, Q^{t_2-1}, ..., Q^{t_{n_T}\text{-}1}, Q^T\}$$

One initializes $\mathcal{R}_{min} = \emptyset$ and $k = card(\mathcal{R}_{min}) = 0$

(2) one identifies the rule $r$ which belongs to the greatest number of subsets from that list. One takes $r$ as a new element of $\mathcal{R}_{min}$ and k is incremented

(3) one removes from the list all the subsets which contain $r$. If the list is now empty, the algorithm is finished, otherwise one goes back to step (2).

##### Results for rat behavioral experiments

We applied the non permissive analysis and the RRS algorithm to the behavioral data recorded from 5 rats and a total of 3322 trials. The following table gives the minimal number of transitions, and the minimal (or approximately minimal) number of rules in RRS, starting with different sets of allowed rules $\mathcal{R}$. We tested 10 sets of rules. The first two sets contain the two cue-independent rules (Left or Right, $L/R$) plus the two single cue dependent rules $L_0$ (rules based on the current light cue) and $M_1$ (rules based on the previous movement). The following seven sets are defined as the full set of 256 rules dependent on the indicated group of three cues. The last set is the union of the seven full sets of 256 rules. Note that this big set contains several instances of the same rule (*e.g.* the Left rule appears in each of the full sets of 256 rules, therefore, it appears seven times in the so-called All set).

| $\mathcal{R}$ | $card(\mathcal{R})$ | $min\, n_T$ | $card(RRS)$ |
|---|---|---|---|
| $L_0 + L/R$ | 4 | 874 | 4 |
| $L_0 + M_1 + L/R$ | 6 | 739 | 6 |
| $L_0M_1R_1$ | 256 | 498 | 43 |
| $L_0L_1R_1$ | 256 | 476 | 49 |
| $L_0L_1M_1$ | 256 | 481 | 45 |
| $L_0M_1M_2$ | 256 | 489 | 45 |
| $L_0R_1R_2$ | 256 | 512 | 60 |
| $M_1M_2R_1$ | 256 | 578 | 37 |
| $M_1R_1R_2$ | 256 | 606 | 43 |
| $All$ | 1792 | 372 | 97 |

Table 1: Non permissive analysis and number of rules in the reduced set for various rule sets.

As a general tendency, increasing the size of the rules set yields fewer transitions.

### 2 The permissive analysis

In this analysis, the Dirac likelihood distribution is replaced by:

$$P(M^t = m \,|\, R^t = r; C^t = c) = \alpha + (1\text{-}2\alpha).(m \wedge r(c))$$

This form of likelihood allows the animal to sometimes perform a movement opposite to the rule he is presumed to apply. The probability to perform such an error is quantified by the parameter $\alpha \ll 1$. One simple way to interpret the parameter $\alpha$ is to relax the deterministic hypothesis (H4): applying a rule $r$ consists to perform the movement $r(c)$ with the probability $1-\alpha$ and to perform the opposite movement with the probability $\alpha$. Another way to interpret the parameter is to relax the well controlled cues hypothesis (H3): the rules are deterministic functions of the known cues plus some unknown cue (or unknown rule), which is generally constant, but can change at some unpredictable trials with a probability $\alpha$ so that the animal change its decision. Anyway, the mathematical formulation remains the same.

The permissive analysis is specified by two parameters ($\varepsilon$ and $\alpha$). At each trial, the transition probability can take one among two values as before; the likelihood can also take one among two values: either $\alpha$ if it is an error trial or $1-\alpha$ if not. Let $n_T$ the number of transitions and $n_E$ the number of errors, the Log posterior probability of the whole sequences of rules is :

$$LP(R) = cste\text{-}n_T Log(\frac{(1\text{-}\varepsilon)(n\text{-}1)}{\varepsilon}) - n_E Log(\frac{1\text{-}\varepsilon}{\varepsilon})$$

The issue is then to find the best compromise between minimizing $n_T$ and minimizing $n_E$. The critical parameter is the *permissive factor* $p$:

$$p = Log(\frac{(1-\varepsilon)(n-1)}{\varepsilon})/Log(\frac{1-\alpha)}{\alpha})$$

The cost $J$ to be minimized is simply defined as:

$$J = n_T - n_E/p$$

with:

$$LP(R) = cste\text{-}J.Log(\frac{(1\text{-}\varepsilon)(n\text{-}1)}{\varepsilon})$$

We first remark that if $p < 0.5$ the permissive analysis turns out to be equivalent to the non permissive analysis, *i.e.* the permissive factor is too low. In fact, let's consider the following sequence or rules given by the non permissive model:

$$aa...aabaa...aa$$

There is two transitions and no error, so that the cost is equal to 2. Consider now the following sequence where the unique $b$ is replaced by an error trial. The new sequence:

$$aa...aaaaa...aa$$

contains no transition and one single error. Its cost is equal to $1/p$. Therefore if $p < 0.5$, the permissive analysis cannot give rise to better sequences than the one produced by the non permissive analysis.

Suppose now that $p > 1$. One can see that a full sequence of $a$ interrupted by two$b$:

$$aa...aabbaa...aa$$

has a higher cost that a sequence containing only "a" (two errors are better than two transitions). More generally, any pairs of errors in a sequence would be suppressed. We consider (but this is debatable) that $p > 1$ is too permissive, and would allow the suppression of transitions which are in fact significant. Therefore, we will restrain ourselves to the weakly permissive analysis where $0.5 < p < 1$. In practice, we choose $p = 2/3$ (*i.e.* the cost of two transitions is equal to the cost of three errors).

As far as we know, there is no algorithm - running in a reasonable time - which is able to build the optimal sequence of rules according to the weakly permissive analysis. We then adopt a greedy forward algorithm as followed

(1) one builds the list of $T$ subsets of compatible rules$\{S^1; ..., S^T\}$, *i.e.*:

$$r \in S^t \Leftrightarrow m^t = r(c^t)$$

(2) Initialize $n_T = n_E = 0$ and $Q^1 = S^1$

(3) For $t = 2$ to $t = T$ compute $Q^t = Q^{t-1} \cap S^t$ and continue while $Q^t \neq \emptyset$

(4) If $Q^t = \emptyset$, $t$ is either a transition trial or an error trial. It is an error trial if and only if $Q^{t-1} \cap S^{t+1} \neq \emptyset$ (that is the trial $t$ can be bypassed). In that case, increment $n_E$ and set $Q^t = Q^{t-1} \cap S^{t+1}$. Otherwise, increment $n_T$ and reset $Q^t = S^t$.

#### Results for rat behavioral experiments

We applied the weakly permissive analysis (WPA) to the same data and same set of rules used with the non permissive analysis (NPA). The cost for NPA is always equal to the number of transitions. The cost for WPA depends on the number of transitions and errors ($WPAn_T$ and $WPAn_E$) with a permissive factor $p = 2/3$.

| $\mathcal{R}$ | $NPACost$ | $WPAcost$ | $WPAn_T$ | $WPAn_E$ |
|---|---|---|---|---|
| $L_0 + L/R$ | 874 | 656.0 | 534 | 153 |
| $L_0 + M_1 + L/R$ | 739 | 694.66 | 662 | 49 |
| $L_0M_1R_1$ | 498 | 480.66 | 472 | 16 |
| $L_0L_1R_1$ | 476 | 466.66 | 458 | 10 |
| $L_0L_1M_1$ | 481 | 471 | 465 | 9 |
| $L_0M_1M_2$ | 489 | 470.33 | 453 | 26 |
| $L_0R_1R_2$ | 512 | 479 | 447 | 48 |
| $M_1M_2R_1$ | 578 | 560.66 | 546 | 22 |
| $M_1R_1R_2$ | 606 | 562.66 | 524 | 58 |
| $All$ | 372 | 369.66 | 367 | 4 |

Table 2: Comparison of the weakly permissive and non permissive analysis for various rule sets.

We remark that the permissive factor benefits mainly to the smallest set of allowed rules. Still, the lowest cost and the minimal number of transitions are obtained with the big All set of rules (which, as we have seen before can be reduced to 97 different rules). In second position, the groups specified by the cues$L_0L_1R_1$,$L_0M_1M_2$ and $L_0L_1M_1$ make quite a good job with only 49 and 45 rules.

Finally, if we look at the most frequently used rule (by counting the number of valid subsets $Q^t$ containing a given rule), we observe that the "persistence rule" (which consists to reproduce the previous movement) is generally ahead, of course providing that it is included in the allowed set $\mathcal{R}$. Within the biggest rule set, the most frequently used rule consists to reproduce the previous movement if it has been rewarded, otherwise to reproduce the movement performed two trials before (the penultimate movement).

#### Incremental Analysis

In this section, we start with only two complementary rules (e.g. Left and Right or Light and Dark or Altern and Persist). Then, we look for the best additional rule which minimizes the NPA cost (i.e. the number of transitions without errors). The process is repeated until no further improvement can be achieved. Because the process is incremental (and not decremental as in the search for Reduced Rule Set) we expect to obtain at the end a NPA cost slightly higher than in the previous analysis. The advantage is to get a (monotonically decreasing) curve for NPA cost as a function of the number of rules.

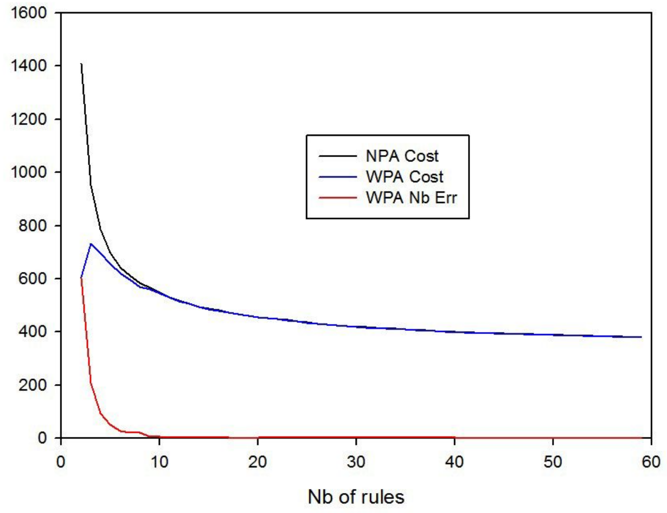


Fig. 1: Minimal cost as a function of the number of rules, obtained by the incremental analysis based on minimal NPA cost, and starting with the two complementary rules Left and Right.

The results depend on the choice of the pair of complementary rules, as indicating in the following table. The influence of the starting pair is mostly at the beginning of the process (*i.e.* for a relatively small number of rules). The best choice appears to be (Altern, Persist).

| Start | 2 | 4 | 6 | 8 | 16 | End | Final Nb |
|---|---|---|---|---|---|---|---|
| L + R | 1409 | 786 | 642 | 586 | 481 | 380 | 59 |
| A + P | 1082 | 742 | 638 | 582 | 478 | 384 | 58 |
| L + D | 1535 | 789 | 672 | 602 | 484 | 383 | 57 |

Table 3: Minimal NPA cost obtained by the incremental analysis starting with the three complementary pairs: L+R = (Left, Right); A+P = (Altern, Persist) L+D = (Light, Dark). The last two columns give the final number of transition (End) and the final number of rules (Final Nb) when the algorithm stops, i.e. when no decreasing cost can be obtained.

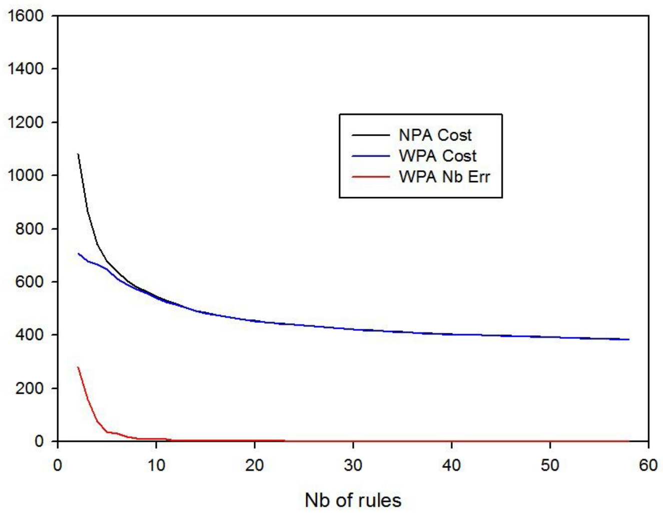


Figure 2: Results of the incremental analysis based on the minimal NPA cost, and starting with the (Altern, Persist) pair.

The same incremental analysis can be used with the minimal WPA cost, in order to obtain this time a monotonically decreasing curve for WPA. As before, one can compare the resulting costs for the three starting pairs of complementary rules (see Table 4).

| Start | Value | 2 | 4 | 6 | 8 | 16 | End | Final Nb |
|---|---|---|---|---|---|---|---|---|
| L + R | Cost | 605 | 584 | 575.7 | 572 | 562.7 | 562.7 | 16 |
| L + R | Errors | 603 | 474 | 472 | 471 | 448 | 448 | 16 |
| A + P | Cost | 708.7 | 655.7 | 588 | 556.3 | 476.7 | 383.7 | 52 |
| A + P | Errors | 280 | 157 | 99 | 59 | 10 | 4 | 52 |
| L + D | Cost | 824.3 | 632.7 | 599.7 | 568.7 | 492.33 | 393.33 | 60 |
| L + D | Errors | 533 | 193 | 115 | 94 | 35 | 8 | 60 |

Table 4: Minimal WPA cost and number of errors obtained by the incremental analysis (based on the WPA criterion) starting with the three complementary pairs.

Note that the Left/Right pair obtains the best initial cost, at the price of a very large number of errors. The cost and the number of errors can be only slightly improved by increasing the number of rules. Again the winner starting pair appears to be Altern/Persist.

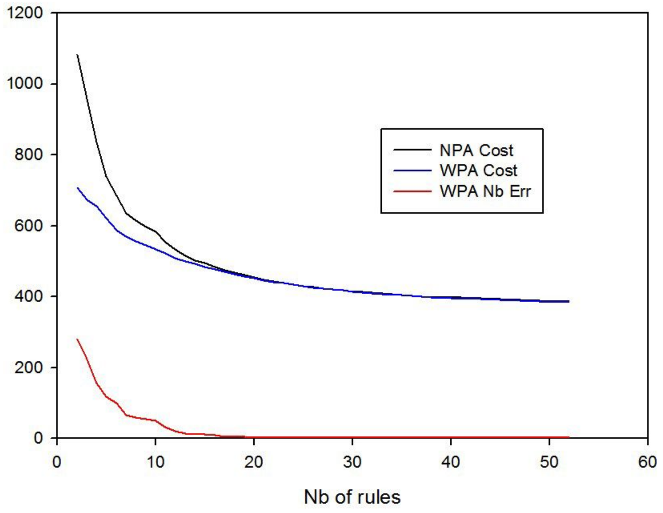


Fig. 3: Results of the incremental analysis based on the WPA cost and starting with the (Altern, Persist) pair.

In general, it is quite hard to understand the "logic" of the first rules which are added to the initial pair by the incremental analysis. To give one single example, starting with the Altern / Persist pair, and looking for the lowest WPA cost, the first two new rules that one of the rat (R20) seems to apply are defined as follows:

- if the previous trial was rewarded, move to the dark arm (*i.e.* $Not(L_0)$); if not previously rewarded, move to the arm which was previously lightened (*i.e.* $L_1$)

- if the previous trial was rewarded, move to the lightened arm ($L_0$); if not previously rewarded, move to the arm which was previously in the dark ($Not(L_1)$)

These two rules are in this case complementary, but this is exceptional. Note that these two rules can explain why the rat seems to have learned the light and dark experimenter's rules.

#### Histograms of sequence lengths

In this analysis, we compute the number of sequences of a given length where the same rule (or the same subset of rules) applies to consecutive trials, for a given set of rules $\mathcal{R}$ and for both NPA and WPA. Below are shown the results for the group of 6 rules (left, right, light, dark, altern & persist), for the group of 256 rules based on the cues $L_0L_1R_1$ and the full set of 1792 rules (All set), according to the WP analysis.

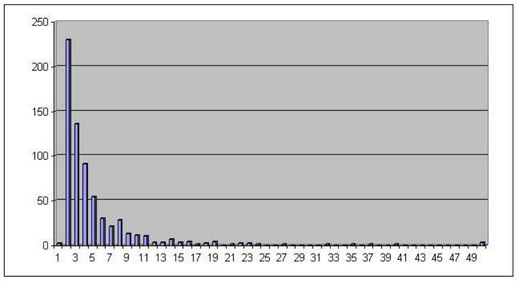
250
200
150
100
50
0
1 3 5 7 9 11 13 15 17 19 21 23 25 27 29 31 33 35 37 39 41 43 45 47 49

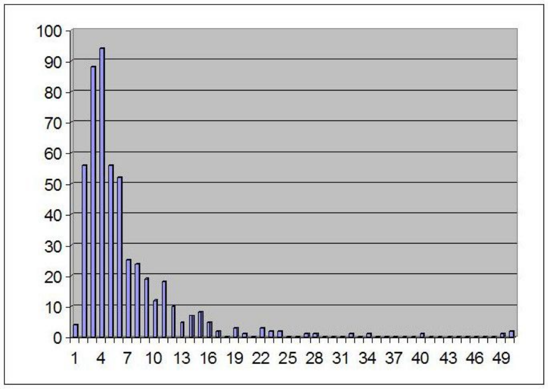
100
90
80
70
60
50
40
30
20
10
0
1 4 7 10 13 16 19 22 25 28 31 34 37 40 43 46 49

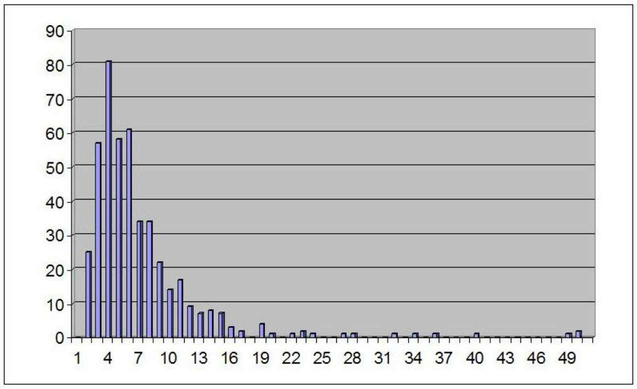
90
80
70
60
50
40
30
20
10
0
1 4 7 10 13 16 19 22 25 28 31 34 37 40 43 46 49

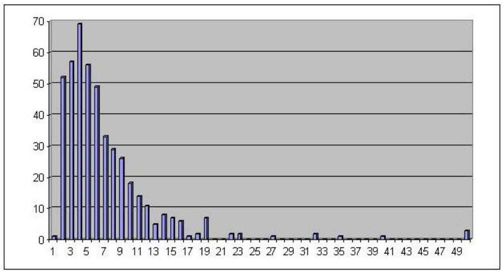
70
60
50
40
30
20
10
0
1 3 5 7 9 11 13 15 17 19 21 23 25 27 29 31 33 35 37 39 41 43 45 47 49

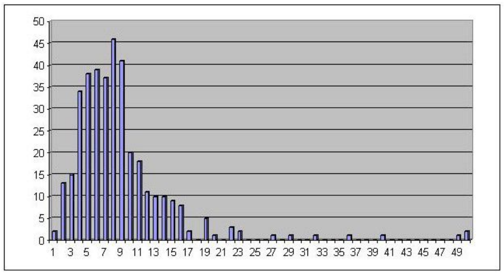


Fig. 4: Histograms of sequence length with WPA applied to various sets of rules. From top to bottom: group of 6 basic rules (left, right, light, dark, altern, persist) with a total of 663 sequences, group of 12 best rules with a total of 496 sequences, group of 20 best rules with a total of 451 sequences, group of 256 rules based on the cues $L_0L_1R_1$ with a total of 459 sequences and finally for the group of 1792 rules with a total of 368 sequences. Note that the peak shifts from 3 to 8, as the number of rules increases.

#### Significance

A quite difficult (because it's ill posed) issue is to evaluate the significance of the NPA or WPA results. One simple way is to compute the probability of any sub-optimal sequence of rules within a given frame, *i.e* within NPA or WPA for a given set of rules. Within NPA, the probability of a sequence of rules is uniquely determined by the number of transitions $n_T$. Within WPA, the probability of a sequence of rules is uniquely determined by the number of transitions $n_T$ and the number of errors $n_E$, though the key number is the cost $J = n_T + n_E/p$. The cost is proportional to $-Log(P)$ with a proportionality coefficient equal to $Log(\frac{(1-\varepsilon)(n-1)}{\varepsilon}) = Log(n-1) + Log(\frac{(1-\varepsilon)}{\varepsilon})$. The parameter $\varepsilon$ has no influence on the selection of the optimal sequence, but it has a key impact on the probability of sub-optimal sequence. In the worst case, we admit $\varepsilon = 0.5$ (which means that the probability to change the rule from trial to trial is 50%). In this worst case, within NPA, a sub-optimal sequence which consists to replace a given sequence $a...a$ by the sequence $a..aba...a$ has the following relative probability (with respect to the optimal sequence $a...a$) function of the number of rules $n$:

| n | p |
|---|---|
| 4 | 0.111 |
| 6 | 0.04 |
| 12 | 0.00826 |
| 20 | 0.00277 |
| 40 | 0.00066 |
| 90 | 0.000126 |

According to the scientific standard in biology, it's significant when $n \geq 6$. Within WPA, one can consider the sub-optimal sequence $a....a$ (no transition, two errors, cost=3) with respect to the optimal sequence *a..abba...a* (two transitions, no error, cost = 2). The relative probability of this sub-optimal sequence is:

| n | p |
|---|---|
| 4 | 0.333 |
| 6 | 0.2 |
| 12 | 0.0909 |
| 20 | 0.0526 |
| 40 | 0.0256 |
| 90 | 0.0112 |

Again, according to the scientific standard, a sub-optimal sequence of rules is significantly sub-optimal for n > 20. One can also compute the relative probability of a sub-optimal sequence *a....a* (no transitions, 3 errors, cost = 9/2) with respect to the optimal sequence *a..abbba...a* (two transitions, no error, cost = 2):

| n | p |
|---|---|
| 4 | 0.06415 |
| 6 | 0.0179 |
| 12 | 0.00249 |
| 20 | 0.000635 |
| 40 | 0.000105 |
| 90 | 0.0000133 |

Clearly, the sub-optimal sequence is significantly (according to scientific standard) less probable as soon as n > 4.

#### Analysis of random movements

Another way to assess the significance of NPA and WPA is to run these methods on a set of (pseudo) random movements. For this, we create pseudo data files with the same number of trials and same cues except for the movement which was randomly drawn with the Mersenne-Twister random generator.

| $\mathcal{R}$ | NPA Cost | WPA Cost | WPA $n_T$ | WPA $n_E$ |
|---|---|---|---|---|
| $L_0 + L/R$ | 1206 | 977.7 | 819 | 238 |
| $L_0 + M_1 + L/R$ | 1061 | 975.0 | 903 | 108 |
| $L_0M_1R_1$ | 678 | 659.7 | 643 | 25 |
| $L_0L_1R_1$ | 697 | 672.3 | 651 | 32 |
| $L_0L_1M_1$ | 620 | 613.0 | 603 | 15 |
| $L_0M_1M_2$ | 636 | 619.7 | 605 | 22 |
| $L_0R_1R_2$ | 674 | 636.7 | 608 | 43 |
| $M_1M_2R_1$ | 706 | 680.66 | 658 | 34 |
| $M_1R_1R_2$ | 720 | 674.0 | 632 | 63 |
| *All* | 448 | 447.3 | 446 | 2 |

Table 5: Comparison of the weakly permissive and non permissive analysis for various rule sets, applied to the random data. There is obviously a dramatic increase of the cost, for both NPA and WPA, and all sets of rules.

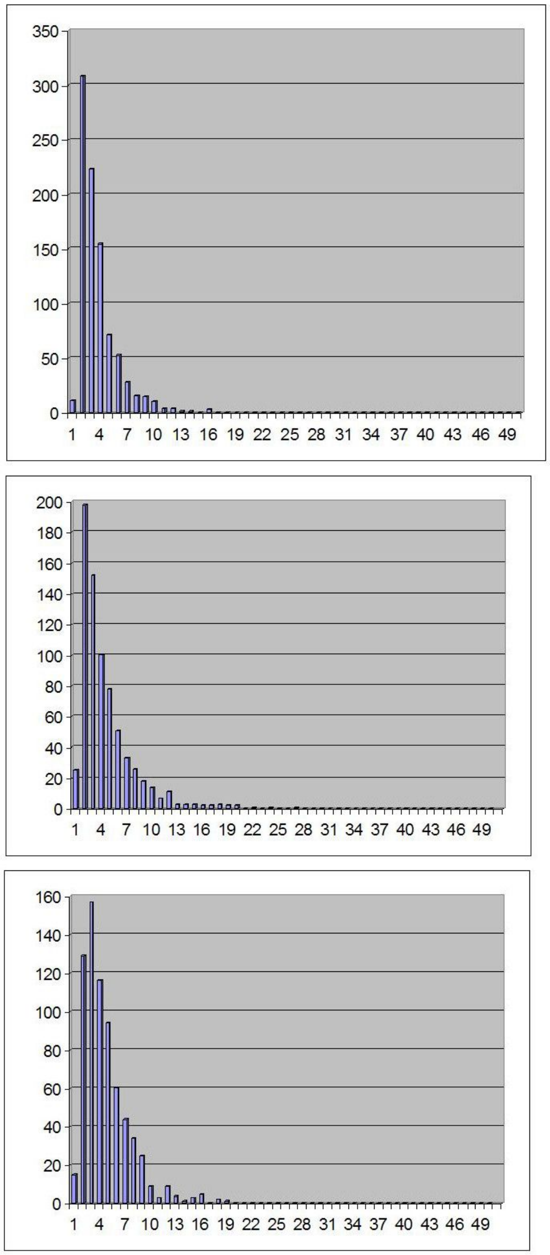

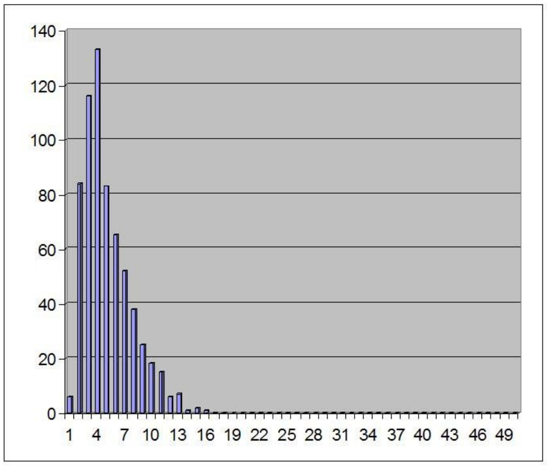


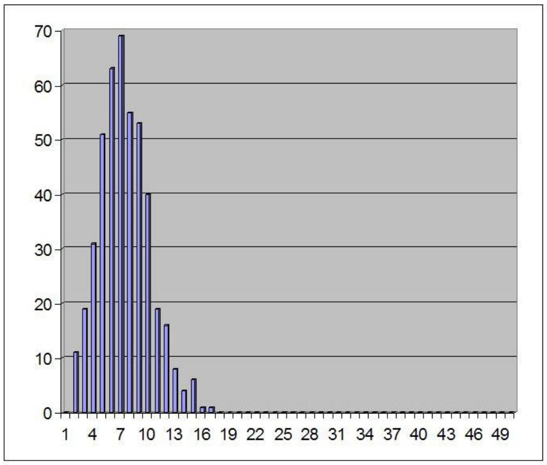


Fig. 5: Histograms of sequence length with WPA applied to the random behavior. The sets of rules are the same than in Fig. 4, *i.e.* from top to bottom: 6, 12, 20, 256 and 1792 rules; the total numbers of sequences are, respectively, 904, 615, 669, 604 and 447. The numbers of errors are also very high, except for the larger sets: 108, 326, 158, 32 and 2. As compared to real data, the peak is sharper and shifts less, as the number of rules increases.

Consider for instance the group of 12 rules (a reasonable size). Compared to random data analysis, the real data analysis provides significantly less short sequences (length $< 4$), roughly similar number of medium size sequences (length between 4 and 12) and a higher number of long sequences (length $> 12$). Similar conclusion can drawn for the sets of 6 and 20 rules. For 6 rules, the real rat results exceeds the random results for sequences longer than 10 (though 3 sequences of length 16 are found in the random data, against 4 for the real rat...). For 20 rules, real rat results exceeds random results for sequences longer than 12, though one sequence of length 19 is found in the random data). As a consequence, one cannot exclude that randomness can account for the existence of one long sequence of consecutive trials (up to about 20 trials). It is the number of long sequences that matters. If we look at the subset of rules which are compatible with sequences of length $> 12$, most rules of the reduced rule set are observed. For instance, in the ascending method 11 among the first 12 best rules, and 15 among the first 20 best rules are observed during long sequences.

#### Outside NPA & WPA: the unknown rule ...

We consider in this section the hypothesis that the rat can use sometimes an "unknown rule" $u$ in addition to the regular rules $r \in \mathcal{R}$. As this unknown rule explains any movement, it should be penalized in some way. Otherwise, the whole data would be perfectly explained by a unique sequence of $u$, without transition and without error. The weak permissive analysis was a first way to introduce this idea; the penalization was quantified by the permissiveness factor. However, this factor was set to a minimal value of 2/3, in order to prevent the possibility of two consecutive errors. Another way to formalize this idea is to penalize $u$ through the specification of transition probability. We need now three parameters in order to define:

- as before the transition probability between two different regular rules $\varepsilon/(n-1)$

- the transition probability from regular to unknown rule: $\varepsilon_1$

- the transition probability from unknown to regular rule: $\varepsilon_2$

The difficulty of this approach results from the arbitrary choice of these parameters. As in section 2, the parameters can be translated into cost *i.e.* the change in Log P resulting from a regular transition $c_t$, a transition towards $u$ $C_u$, a return transition from $u$ $C_r$ and finally the cost to maintain $u$ between two consecutive trials $C_m$. Simple reasoning shows that we must have :

(1) $C_u + C_r < 2C_t$ otherwise $u$ will never be used to account for the data

(2) $C_u + C_r + kC_m > 2C_t$ for some maximal length $k$ otherwise infinite sequence of $u$ will be used instead of regular rules. These considerations lead to some restriction for the parameters, but they cannot be precisely specified without some arbitrary choice.
